# Chemical Inhibitors of DksA1, a Conserved Bacterial Transcriptional Regulator, Suppressed Quorum Sensing-Mediated Virulence in *Pseudomonas aeruginosa*

**DOI:** 10.1101/2020.12.10.420422

**Authors:** Kyung Bae Min, Wontae Hwang, Kang-Mu Lee, Sang Sun Yoon

**Author notes:** Corresponding author, Sang Sun Yoon, Ph.D., Department of Microbiology, Yonsei University College of Medicine, 250 Seongsanno, Seodaemun-gu, Seoul, 120-752, Korea.

## Abstract

*Pseudomonas aeruginosa* is a Gram-negative bacterium of clinical importance. As an opportunistic pathogen, its virulence is dependent on quorum sensing (QS). Stringent response (SR), a conserved mechanism in the bacterial kingdom, is activated under diverse stress conditions including nutrient starvation. DksA1, an RNA polymerase-binding transcriptional regulator, plays a role in *P. aeruginosa* SR. Our recent study clearly demonstrated that, apart from SR, DksA1 regulates a wide range of phenotypes including QS-mediated virulence. This suggests that DksA1 is a potential target to be inhibited to control *P. aeruginosa* infections. Here, we screened a library of 6,970 chemical compounds and identified two compounds (termed Dkstatin-1 and Dkstatin-2) that specifically suppress DksA1 activity. Two Dkstatins substantially suppressed production of elastase and pyocyanin and protected a murine host against lethal infection with PAO1, a prototype strain of *P. aeruginosa*. Dkstatins also suppressed the production of homoserine lactone (HSL)-based autoinducers that activate *P. aeruginosa* QS. The level of 3-oxo-C12-HSL produced in Dkstatin-treated cells was almost identical to that in the Δ*dksA1* mutant of PAO1. Our RNASeq analysis showed that transcript levels of genes involved in QS and virulence were markedly reduced in Dkstatin-treated PAO1 cells, further demonstrating that Dkstatin-mediated suppression occurs at the transcriptional level. Importantly, Dkstatins increased antibiotic susceptibilities of PAO1, particularly to the antibiotics that inhibit protein synthesis. Our co-immunoprecipitation assay demonstrated that Dkstatins interfere with the binding of DksA1 to the β subunit of RNA polymerase, which potentially explains the mode of Dkstatin action. Collectively, our results suggest that the inhibition of *P. aeruginosa* QS, which has often been attempted, can be achieved by DksA1 inhibitors. Dkstatins will prove to be useful when establishing infection control strategies.

**Author summary:** Bacterial cells developed numerous systems to handle environmental stresses. Stringent response (SR) is one of those systems, and it is activated under the stress of nutrient starvation. Active SR requires DksA1, a protein conserved across bacterial species. Besides this well-recognized function, DksA1 in *P. aeruginosa* positively regulates its virulence by activating quorum sensing (QS). This finding led us to hypothesize that inhibiting DksA1 would result in virulence attenuation of the pathogenic *P. aeruginosa*. Herein, we identified two molecules that effectively suppressed DksA1 function through a large-scale compound library screen. These two compounds suppressed *P. aeruginosa* virulence and therefore protected a murine host from a deadly infection of *P. aeruginosa*. Moreover, the compounds also rendered *P. aeruginosa* more susceptible to antibiotics. Our results demonstrate that dangerous *P. aeruginosa* infections can be controlled by inhibiting DksA1.

## Introduction

*Pseudomonas aeruginosa*, an opportunistic human pathogen, has extensive metabolic capabilities of adapting to diverse environments including immunocompromised human hosts [1]. In addition, *P. aeruginosa* commonly contains high proportions of regulatory genes, particularly those for diverse signal pathways that establish resistant phenotypes [1, 2]. Stringent response (SR) is a highly conserved mechanism across bacterial species, activated in response to nutrient starvation [3]. SR is mediated by two key elements, nucleotide alarmones called guanosine tetra- and penta-phosphate, (p)ppGpp, and a transcriptional regulator DksA [4, 5]. DksA is a 17 kDa protein with a coiled-coil N-terminal domain and globular C-terminal domain consisting of a Zn^2+^ binding motif with α-helix structures [3, 6]. According to the structural analysis, the Zn^2+^ binding motif of DksA consists of four cysteine residues, which play a key role in sustaining the folding of the C-terminal and coiled-coil regions of DksA [7]. Under nutrient starvation, a RelA enzyme is introduced to tRNA for the purpose of sensing amino acid deficiency and initiating the synthesis of (p)ppGpp via GTP and GDP consumption [5, 8]. Using (p)ppGpp, DksA binds to RNA polymerase (RNAP) for downstream transcriptional regulation, such as the repression of rRNA gene transcription [3, 5, 9].

The mode of action regarding the interaction of DksA with RNAP was uncovered in a series of studies using *E. coli*. In *E. coli*, the intracellular level of DksA remains constant throughout the lifecycle, unlike (p)ppGpp [10]. The constant level of DksA was initially considered to be crucial as it was identified to suppress *dnaK* mutation with its chaperon activity [3, 10]. However, DksA was later revealed to be even more significant as it serves as a transcriptional suppressor of rRNA and ribosomal proteins in *E. coli* [3, 8]. DksA directly binds to RNAP and modulates RNAP activity by destabilizing the open complex to prevent intermediate complexation by competition for transcription initiation [3, 4, 11]. A current model demonstrates that DksA binding requires multiple interactions with (i) rim helices of the β’-subunit, (ii) an active site of the β-subunit and (iii) a β-subunit site insertion 1 (β-SI1) in a secondary channel of RNAP [11].

DksA is also critically involved in regulating bacterial pathogenesis in several pathogens [10, 12–15]. In *Vibrio cholerae*, DksA upregulated expression of *fliA*, encoding a sigma factor that regulates its motility. Furthermore, uninterrupted production of cholera toxin and hemagglutinin protease required functional DksA [16]. Likewise, in *Salmonella*, DksA was found essential for the expression of motility, biofilm formation, cellular invasion and intestinal colonization that caused *in vivo* systemic infection [17]. Moreover, DksA in *Salmonella* controlled central metabolism to balance its redox state, which in turn helped resist against oxidative stress produced by anti-microbial phagocytes [18].

*P. aeruginosa* harbors five genes in its genome encoding proteins belonging to the DksA superfamily, including two that are highly homologous to the typical DksA in *E. coli* [12, 19]. Of these two DksA homologs, named DksA1 and DksA2, DksA1 is structurally and functionally similar to *E. coli* DksA. DksA2, on the other hand, was reported to only partially replace DksA1 functions, as it lacks the typical Zn^2+^ binding motif present in DksA [19]. However, our recent study clearly suggested that DksA1, not DksA2, plays a dominant role as a suppressor of ribosomal gene expression [13]. Importantly, a Δ*dksA2* mutant exhibited almost identical phenotypes with its parental strain, PAO1, indicating that DksA2 can be dispensable. Beyond its traditional function, DksA1 was also identified to regulate a wide range of phenotypes including quorum sensing (QS)-related virulence, anaerobiosis and motilities [13]. Based upon these findings, we hypothesized that DksA1 may be an efficient target for inhibiting *P. aeruginosa* infection.

In the present work, we screened a library of chemical compounds (n=6,970) and identified two molecules that effectively compromised DksA1 activity. PAO1 cells treated with each candidate compound shared much of the characteristics of the Δ*dksA1* mutant, such as significant attenuation of QS-mediated virulence and elevated antibiotic susceptibility. Given that QS machinery has been a target for inhibition, our results demonstrate that DksA1 can serve as a novel avenue to achieve *P. aeruginosa* QS inhibition.

## Results

### Screening a library of chemical compounds for DksA1 inhibitors

To set up a screening scheme in a high-throughput manner, we needed to find a phenotype of the Δ*dksA1* mutant that can be easily and reproducibly measured. In Salmonella, it was reported that DksA modulates the redox balance in response to oxidative stress, and the NADH level decreased when the *dksA* gene is disrupted [18]. We therefore examined whether the phenotype observed in Salmonella is also detected in *P aeruginosa*. NADH levels are indirectly represented by the conversion of tetrazolium salt to formazan (purple pigment), which is mediated by a NADH-dependent reductase [20]. In accordance with what was observed in *Salmonella*, the *P aeruginosa* Δ*dksA1* mutant produced a noticeably reduced amount of formazan (S1A Fig). Since the formazan assay can easily be performed, we concluded that this assay can serve as an efficient screening platform, minimizing the chance of plate-to-plate variations. The library that we used was a compound library consisting of a total of 6,970 chemicals that was provided by the Korean Chemical Bank. The screening was conducted in two stages. To begin with, we screened the entire library for the formazan assay to isolate candidate compounds. Then, we compared phenotypes induced in PAO1 by each candidate compound with those of the Δ*dksA1* mutant (S1B Fig). Our goal was to isolate compounds that make PAO1 behave like the Δ*dksA1* mutant. In the first step, we screened out a total of 178 chemical compounds including 25 compounds that robustly reduced formazan production and 153 mild reducers. In a repetitive formazan assay using these 153 mild reducers, we selected another set of 25 compounds that exhibited relatively strong activities in reducing formazan production. Consequently, a total of 50 compounds were selected in the first stage (S1B Fig).

It was previously reported that transcription of genes encoding ribosomal proteins is highly activated in the Δ*dksA1* mutant [12, 13]. We postulated that these marked changes at the transcriptional levels may allow us to verify the phenotypes induced by candidate compounds. We therefore constructed a reporter *P. aeruginosa* strain by chromosomally integrating an *rpsB* gene promoter fused with *lacZ* ORF (P_*rpsB*_::*lacZ*). The *rpsB* gene codes for 30S ribosomal protein S2. In our previous work, we also reported that the production of elastase, a dominant virulence determinant and an easily measurable phenotype, is highly suppressed in the Δ*dksA1* mutant [13]. Therefore, we also assessed elastase production as a second-stage verification. By successive β-galactosidase and elastase activity assays, we finally nominated a total of 4 chemicals, 55B05, 02G09, 86B09 and 45G08, as a set of final candidates. We then monitored these two phenotypes in diverse cultivation conditions and at different time points to finally select chemicals that yield consistent results. While reduced elastase production and elevated *rpsB* transcription were observed at 4 hour post-inoculation by all 4 chemicals at a 50 *μ*M concentration, the effects of 02G09 and 45G08 did not last for the extended period of incubation (S2A and S2B Fig). At 8 hour post-inoculation, the levels of elastase produced in 02G09- or 45G08-treated PAO1 were almost identical to that of PAO1 (S2B Fig). Thus, 55B05 and 86B09 were finally selected as DksA1 inhibitors and we referred to these as Dkstatin-1 and Dkstatin-2.

### Structure and specificity of Dkstatin-1 and Dkstatin-2

The Korean Chemical Bank database provides the structures of Dkstatin-1 and Dkstatin-2, and they were found to have distinct chemical structures (Fig 1A and 1B). To further confirm that Dkstatin molecules act specifically against DksA1 activity, formazan production and *rpsB* reporter expression were measured in PAO1 and the Δ*dksA1* mutant as well. Hereafter, Dkstatin-1 and −2 were used at 150 *μ*M concentrations in all experiments. Again, levels of formazan production in PAO1 were noticeably reduced in the presence of Dkstatin treatment (Fig 1C). It is of interest that formazan production in PAO1 treated with 150 *μ*M Dkstatin-1 was even lower than that of the Δ*dksA1* mutant. On the other hand, levels of formazan production in the Δ*dksA1* mutant were not further decreased by either Dkstatin compound (Fig 1C). Likewise, Dkstatin-1 and Dkstatin-2 did not further increase the *rpsB* gene transcription in the Δ*dksA1* mutant (Fig 1D). As Dkstatin-1 and Dkstatin-2 were identified to inhibit DksA1 activity, no apparent effects of Dkstatins were present in a strain devoid of *dksA1* gene. These results suggest that Dkstatin compounds specifically and selectively inhibit DksA1 activity.

**Fig 1.**
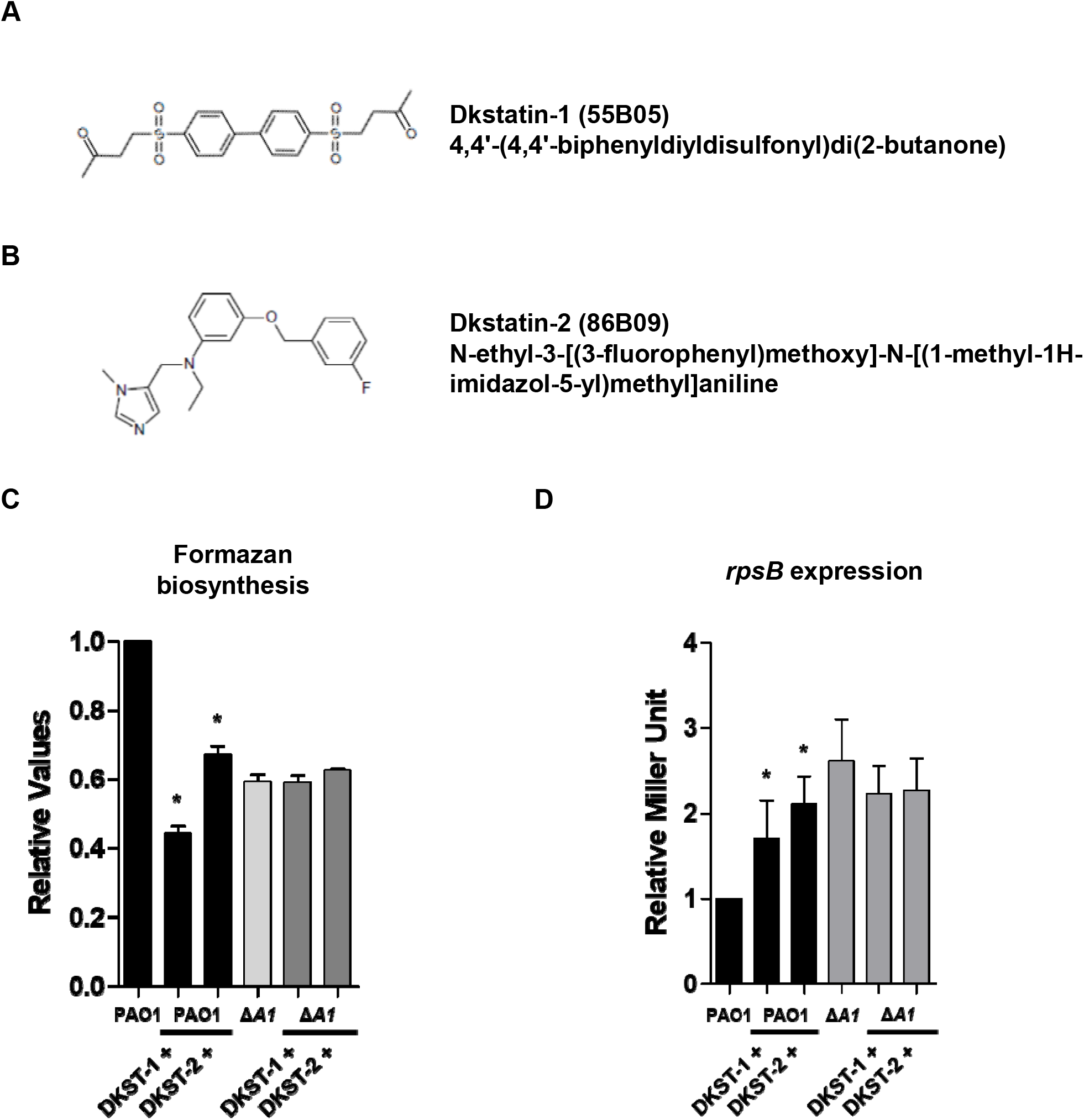
Effect of Dkstatin candidate compounds on *P. aeruginosa*. **(A)** and **(B)** Chemical structure of Dkstatin-1 (DKST-1) and Dkstatin-2 (DKST-2). **(C)** Relative formazan production of PAO1 and Δ*dksA1* mutant grown in LB with 150 μM Dkstatin-1 and Dkstatin-2 for 8 hours. The values of mean ± S.D. are presented (n=3). **(D)** Relative *rpsB* expression of PAO1 P_*rpsB*_::*lacZ* and Δ*dksA1* P_*rpsB*_::*lacZ* strains incubated in LB with 150 μM Dkstatin-1 and Dkstatin-2 for 8 hours. The *rpsB* gene expression is represented as β-galactosidase activity. The values of mean ± S.D. are presented (n=3).

### Dkstatin compounds interrupt QS-mediated virulence in *P. aeruginosa*

DksA1 plays versatile roles at the transcriptional level affecting various phenotypes [13]. Therefore, Dkstatin molecules inhibiting the activity of DksA1 may induce diverse physiological changes in *P. aeruginosa*. To identify whether Dkstatins affect the growth of PAO1, we conducted growth curve experiments of PAO1 in the presence of 150 *μ*M of Dkstatins. Control growth curves were obtained by adding equal volume of DMSO used as a solvent. In comparison with PAO1 and Δ*dksA1* mutant, PAO1 growth never changed upon Dkstatin-1 supplementation (Fig 2A). To our surprise, a noticeable increase in PAO1 growth was repeatedly observed when treated with Dkstatin-2 (Fig 2A). When CFU was enumerated 8 hour post-inoculation, no obvious growth enhancement was observed (Fig 2B). Together, these two sets of results indicate that Dkstatin-1 and −2 at least do not down-regulate bacterial growth and viability.

**Fig 2.**
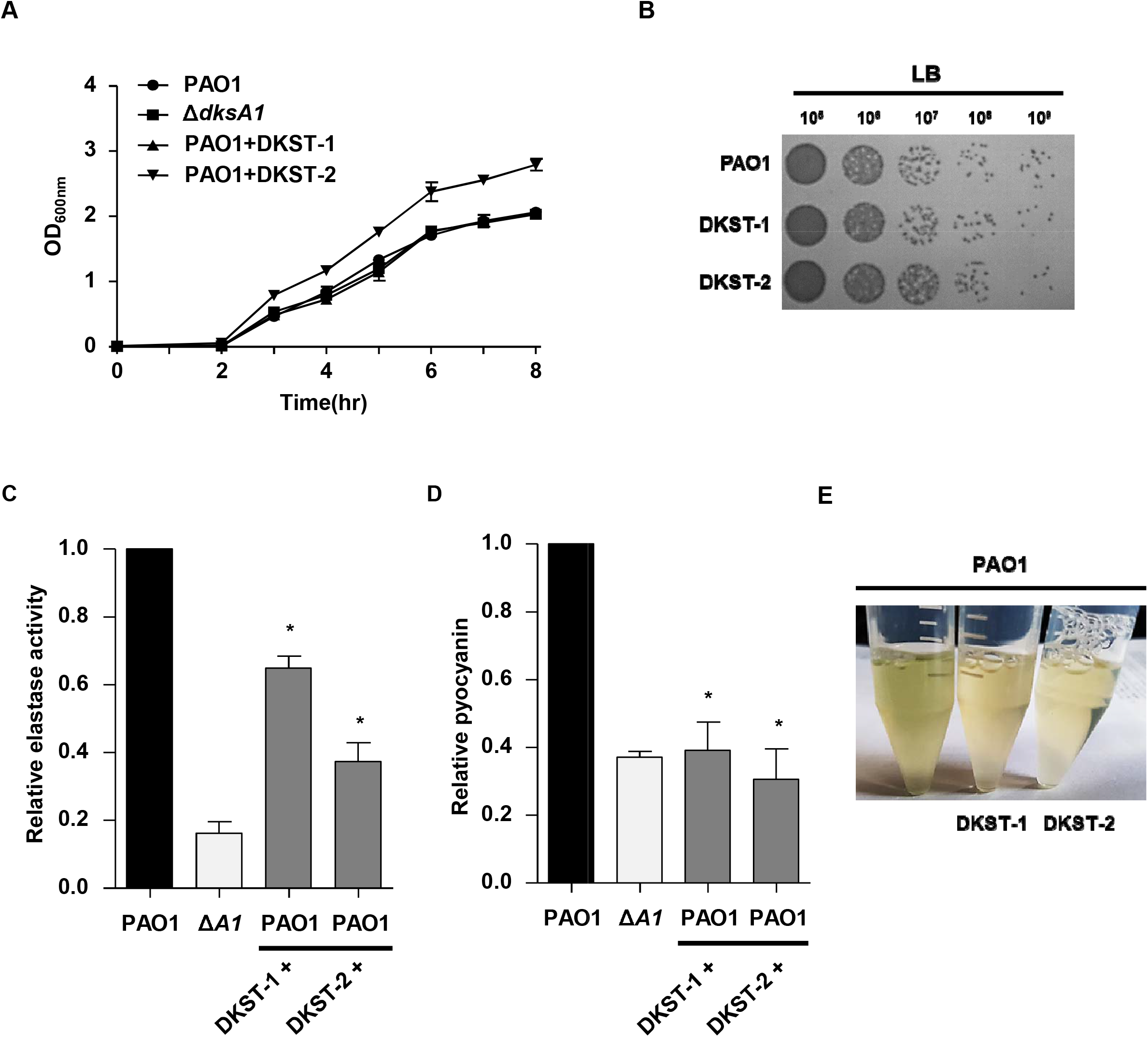
Effects of Dkstatin-1 and Dkstatin-2 on bacterial growth and virulence. **(A)** Growth curves of PAO1 and Δ*dksA1* mutant in plain LB and LB supplemented with 150 μM of Dkstain-1 (DKST-1) or Dkstatin-2 (DKST-2). Aliquots of bacterial cultures (n=3) were withdrawn every hour to measure OD_600_ values. **(B)** Cell viability of PAO1 incubated with Dkstatin-1 (DKST-1) and Dkstatin-2 (DKST-2). **(C)** Relative elastase activity of tested strains. Elastase activity was measured as described in the Methods and was normalized with that of PAO1. The values of mean ± S.D. are presented (n=5). **(D)** Pyocyanin production in each condition was quantified and normalized with that of PAO1. The values of mean ± S.D. are presented (n=3). **(E)** Visual comparison of bacterial culture supernatants. Loss of pigment was clearly observed in PAO1 and both Dkstatin compounds.

At a concentration of 150 *μ*M, Dkstatin-2 suppressed elastase production to a more significant degree (Fig 2C). Under Dkstatin-2 treated conditions, the level of elastase production was less than 40% of what control PAO1 cells produced. Elastase production in PAO1 treated with 150 *μ*M Dkstatin-1 was approximately 65% of the control level. Along with elastase, pyocyanin is another crucial virulence determinant whose production is also controlled by QS in *P. aeruginosa* [2, 21, 22]. When we measured the pyocyanin production, a marked reduction was observed in Dkstatin-treated PAO1 (Fig 2D). Interestingly, the degree of reduction by either Dkstatin molecule in PAO1 was comparable to that induced by the *dksA1* gene deletion (Fig 2D). The loss of pyocyanin production was clearly represented in the image of culture supernatants shown in Fig 2E.

In *P. aeruginosa*, QS is operated by small organic molecules, termed autoinducers [23]. Since major QS-mediated phenotypes were suppressed in Dkstatin-treated PAO1 and in the Δ*dksA1* mutant, we monitored how many autoinducers were produced in response to the Dkstatin treatment. Two homoserine lactone (HSL) autoinducers, 3-oxo-C12-HSL and C4-HSL, were semi-quantitatively measured as described in the Methods section. Based on our quantification, levels of 3-oxo-C12-HSL produced in either Dkstatin-1 or Dkstatin-2 treated PAO1 cells were almost identical to that in the Δ*dksA1* mutant (Fig 3A). Similarly, levels of C4-HSL produced upon treatment with either Dkstatin were noticeably decreased to 53% and 63% of what was detected in the control group, respectively (Fig 3B).

**Fig 3.**
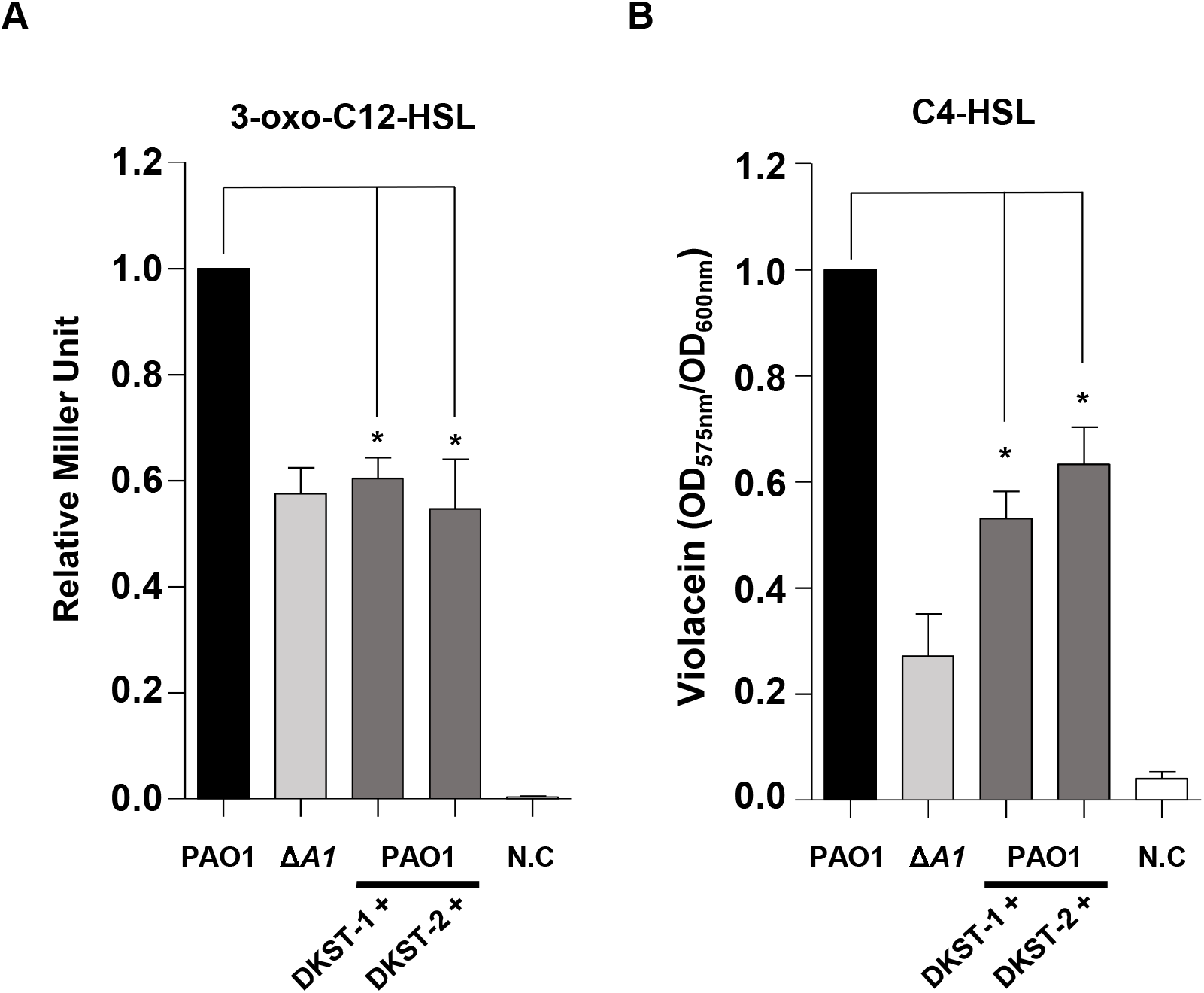
Measurement of 3-oxo-C12-HSL and C4-HSL produced when supplemented with Dkstatin compounds. **(A)** Relative levels of 3-oxo-C12-HSL (C12-HSL) were measured using an *E. coli* reporter strain harboring a pKDT17 plasmid. Concentrations of C12-HSL are represented as β-galactosidase activity. The values of mean ± S.D. are presented (n=5). **(B)** Relative levels of *N*-butyryl-homoserine lactone (C4-HSL) were monitored by measuring the violacein production from the biosensor strain *C. violaceum* CV026. Violacein production was measured at 590 nm absorbance and was normalized to the OD_600_ values of CV026. 150 μM Dkstain-1 (DKST-1) and Dkstatin-2 (DKST-2) were supplemented in all experiments.

When Dkstatin-treated PAO1 cells were co-treated with extraneous C4-HSL and 3-oxo-C12-HSL, elastase production was restored (S3 Fig). Elastase production was almost completely recovered following 4 hour autoinducer complementation, while the complementation effects diminished thereafter (S3 Fig). These results demonstrate that the LasI-R and RhlI-R QS circuits can be substantially impaired by two Dkstatins, with the LasI-R system being suppressed almost to the degree of the Δ*dksA1* mutant. Furthermore, our results indicate that Dkstatin-induced virulence attenuation occurred at the level of the HSL-based autoinducer production.

### Anaerobic respiration is affected by Dkstatins

The transcription of genes involved in denitrification or anaerobic respiration requires DksA1 activity [13]. We therefore examined whether the Dkstatin compounds also have an impact on anaerobic respiratory growth of *P. aeruginosa*. We tested bacterial growth using NO_3_^-^ as an alternative electron acceptor. When cells were grown anaerobically with 25 mM NO_3_^-^, PAO1 strains reached an average OD_600_ value of ~1.6 in 24 hours (Fig 4A). In contrast, final OD_600_ values plateaued at ~1.24 in the Δ*dksA1* mutant under the same conditions. Under the Dkstatin-1 supplemented growth conditions, PAO1 grew up to OD_600_ values of ~1.18, similar to the growth of a Δ*dksA1* mutant. Likewise, average OD_600_ values were ~1.27 in the presence of Dkstatin-2 (Fig 4A). Interestingly, the growth stimulating effect of Dkstatin-2 (Fig 2A) observed during aerobic growth was not detected under anaerobic respiration. In a CFU counting assay, we observed that the addition of Dkstatin-2 resulted in a similar outcome as the Δ*dksA1* mutant (Fig 4B). Around 10-fold decreases in viable cell numbers were observed at the end of the growth experiments both in PAO1 treated with Dkstatin-2 and in the Δ*dksA1* mutant, whereas a significant difference in cell numbers was not detected when Dkstatin-1 was supplemented. Together, these results illustrate that Dkstatin compounds clearly have an impact on DksA1 activity that is involved in activating major anaerobic respiration pathways in *P. aeruginosa*, although the degree of impact varies between the two Dkstatins.

**Fig 4.**
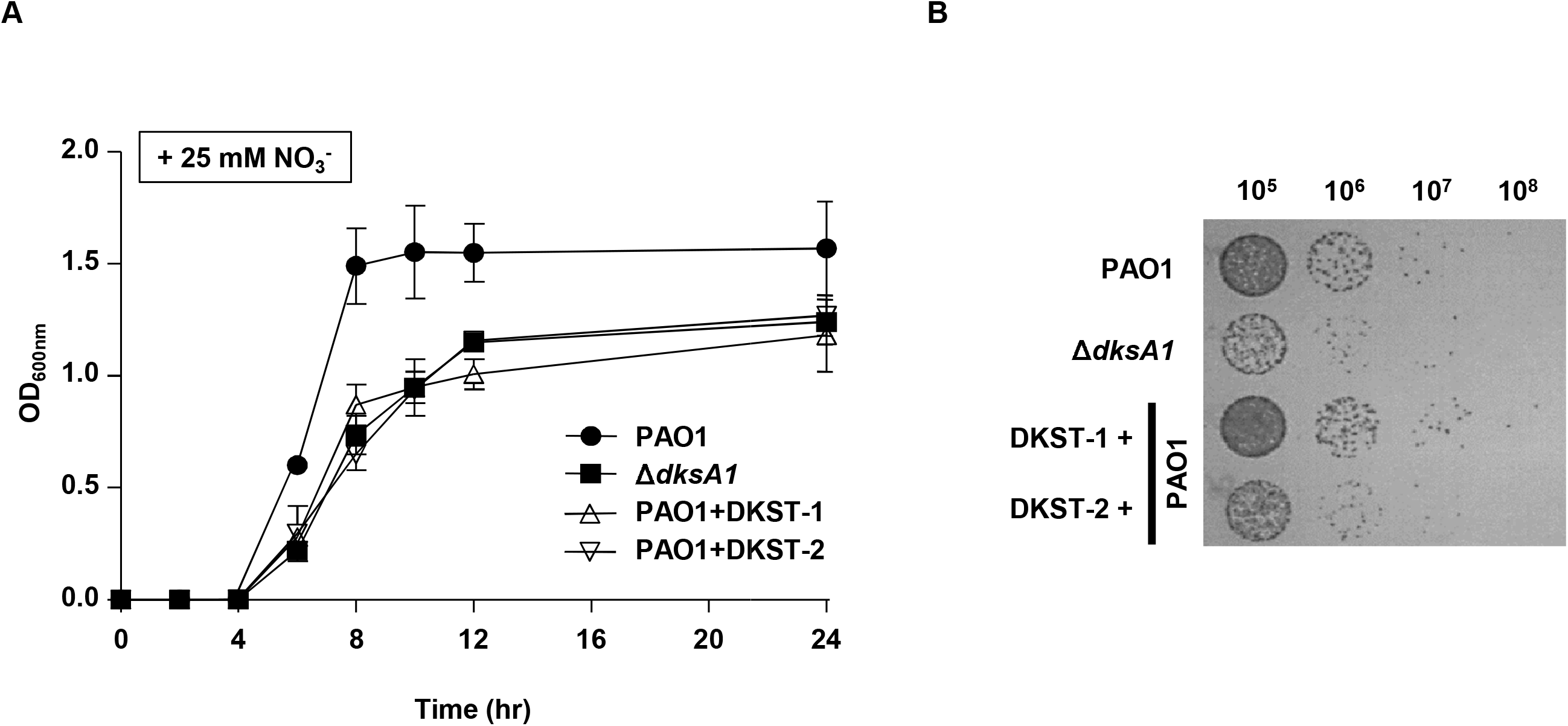
Effects of Dkstatin-1 and Dkstatin-2 on anaerobic respiration. **(A)** Growth curves of PAO1 and Δ*dksA1* mutant in LB and LB supplemented with 150 μM Dkstain-1 (DKST-1) and Dkstatin-2 (DKST-2). 25 mM NO^3-^ was added as an alternative electron receptor. OD values of bacterial culture aliquots (n=2) were measured every 2 hours. **(B)** The cell viability of anaerobically grown Δ*dksA1* mutant and PAO1incubated with 150 μM Dkstain-1 and Dkstatin-2. Bacterial cultures grown anaerobically for 24 hours in LB supplemented with 25 mM NO^3-^ were serially diluted. 10 μl of each diluent was spot inoculated on an LB agar plate. The plates were incubated in anaerobic conditions at 37°C for 24 hours.

### Dkstatin-1 and Dkstatin-2 increase antibiotic susceptibility

*P. aeruginosa* is notorious for its antibiotic resistance [1]. As an opportunistic pathogen, it has also acquired resistance to multiple drugs [1, 24]. Our previous data showed that the Δ*dksA1* mutant became hyper-susceptible to gentamycin or tetracycline at sub-MIC conditions [13]. In contrast, the mutant was not more susceptible to β-lactam antibiotics, such as ampicillin and carbenicillin, than PAO1 [13]. To examine whether the Dkstatin treatment would lead to the interruption of DksA1 activity resulting in increased antibiotic susceptibility, we tested imipenem (Imp), gentamycin (GM), tetracycline (TC), kanamycin (KM), streptomycin (SM) and tobramycin (TB). Dkstatins failed to increase PAO1 susceptibility in response to the treatment with Imp at a 2.5 *μ*g/ml concentration (Fig 5A). Supplementation of Dksatin-1 or Dkstatin-2 both at 150 μM concentration significantly reduced cell viability when PAO1 cells were incubated with 2.5 μg/ml of GM (Fig 5B). Here, a more than 10-fold decrease in cell viability was detected in the presence of either Dkstatin. Interestingly, increased susceptibility by Dkstatin was also observed during incubation with other aminoglycoside antibiotics such as KM (Fig 5D), SM (Fig 5E) or TB (Fig 5F). In the presence of Dkstatin, viable cell numbers of KM-treated PAO1 cells decreased around 10-fold when compared with the DMSO-treated PAO1 control (Fig 5D). Severe loss of viability was observed in PAO1 cells being co-treated with SM and Dkstatin as well (Fig 5E). We also observed that Dkstatin treatment increased PAO1’s susceptibility to 8 μg/ml of TC, an antibiotic that also inhibits protein synthesis machinery (Fig 5C). These results indicate that Dkstatins specifically increase bacterial susceptibilities to antibiotics that specifically target protein biosynthesis in *P. aeruginosa*.

**Fig 5.**
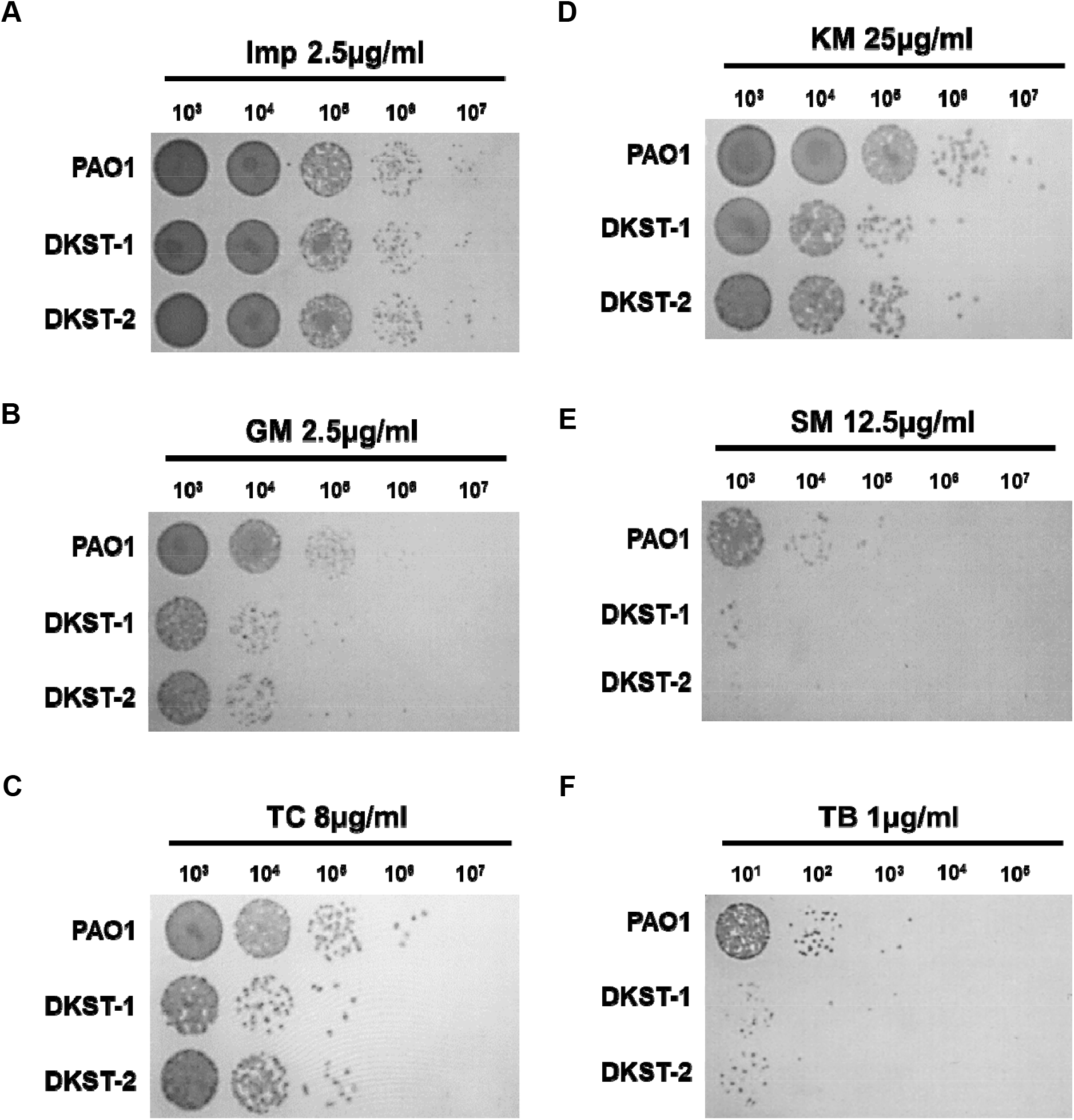
Effects of Dkstatin-1 and Dkstatin-2 on antibiotic susceptibility. CFU counting of PAO1 incubated with Dkstatin-1 (DKST-1) and Dkstatin-2 (DKST-2) when supplemented with antibiotics; (**A**) 2.5 μg/ml of imipenem, (**B**) 2.5 μg/ml of gentamicin (**C**) 8 μg/ml of tetracycline, (**D**) 25 μg/ml of kanamycin, (**E**) 12.5 μg/ml of streptomycin, and (**F**) 1 μg/ml of tobramycin.

### Dkstatins interfere with DksA1’s interaction with RNA polymerase

Action of DksA1 is dependent on its interaction with RNA polymerase (RNAP). It was reported that DksA1 binds to the β subunit of RNAP in *E. coli* [11]. We hypothesized that Dkstatins may work by one of two possible scenarios. First, Dkstatins may inhibit the production of DksA1 in *P. aeruginosa*. Second, Dkstatins may interfere with DksA1’s binding to RNAP. To precisely understand how Dkstatins inhibit DksA1 activity, we constructed a modified PAO1 strain, where the 3’-end of the *dksA1* gene is fused with the nucleotide sequences coding for the 3XFLAG tag. In this strain, DksA1 is translated as a fusion protein with the 3XFLAG tag at its C-terminus. To begin with, we tested whether Dkstatin compounds affect production of DksA1 through western blotting. As a result, western blotting using an anti-FLAG antibody indicates that the level of DksA1 production was never changed by the treatment with either Dkstatin (Fig 6A). The DksA1-specific band was not detected in the sample prepared from the original PAO1 strain (Fig 6A, far-left lane). Next, we conducted a co-immunoprecipitation assay using a magnetic bead (on which an anti-FLAG antibody is immobilized) to examine the effect of Dkstatin treatment on the interaction of DksA1 with RNAP. When precipitated fractions in each preparation were probed with an anti-RpoB antibody, the RpoB band intensity was markedly reduced in both Dkstatin-treated samples, with Dkstatin-2 being superior to Dkstatin-1 in conferring such effects (Fig 6B). Taken together, these results demonstrate that Dkstatins likely act as inhibitors at the post-translational level by interfering with DksA1’s binding to RNAP.

**Fig 6.**
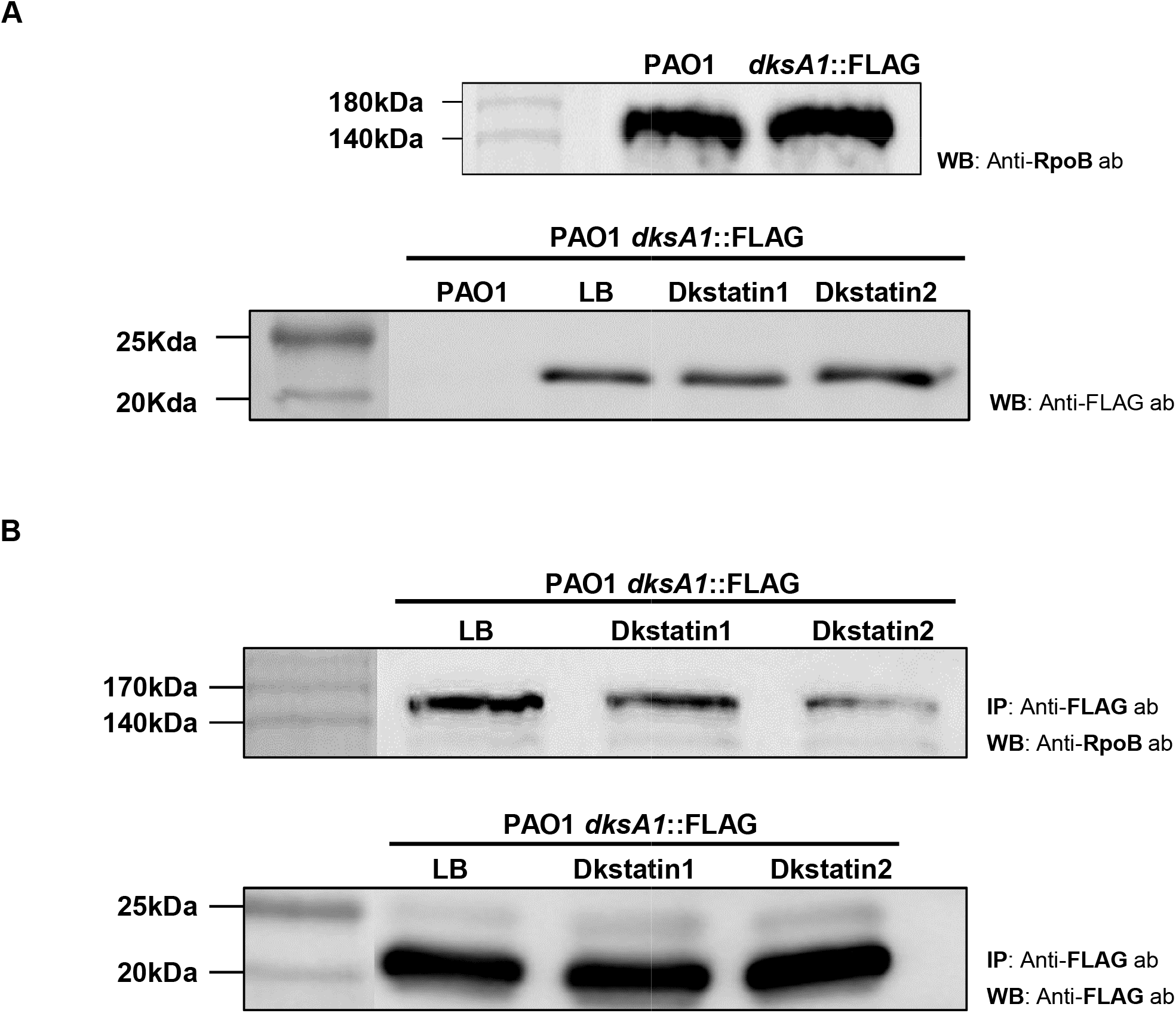
Intracellular production of DksA1 in response to Dkstatin treatment and effects of Dkstatin treatment on binding of DksA1 with RNA polymerase β subunit. (**A**) Original strain of PAO1(first lane) and PAO1 harboring a *dksA1*::FLAG translational fusion construct (following lane) were grown in LB in the presence of either 150 *μ*M Dkstatin-1 (DKST-1) or Dkstatin-2 (DKST-2) for 8 hours. Intracellular fractions were subjected to western blot with an anti-FLAG antibody. As a control, strains were treated with equal volumes of DMSO (Con). Samples at equal protein concentrations were loaded into 12% SDS-PAGE gel. (**B**) A PAO1 strain with *dksA1*::FLAG translational fusion construct was grown for 8 hours in the presence of 150 *μ*M Dkstatin-1 (DKST-1), Dkstatin-2 (DKST-2) or an equal volume of DMSO (Con). Each intracellular extract was incubated with an equal volume of magnetic beads conjugated with an anti-FLAG antibody and were precipitated. Resuspended precipitates were loaded onto SDS-PAGE gel for western blot with the anti-FLAG antibody and the anti-RpoB antibody.

### Dkstatin-1 and Dkstatin-2 induced distinct changes in transcriptome profiles of PAO1

Although two Dkstatins invariably suppressed the activity of DksA1, inducing similar phenotypic changes, the chemical structures of the two molecules are quite different (Fig 1A and 1B). To better understand the effects of Dkstatin treatment, we compared transcriptome landscapes of PAO1 grown without or with either Dkstatin. A global view of the entire transcriptome uncovered the substantial differences between cells treated with Dkstatin-1 or Dkstatin-2 (S4A Fig). This finding was also supported by our PCA plot, which indicates that either treatment induced divergent changes (S4B Fig). As far as the number of affected genes was concerned, the effect of Dkstatin-1 treatment appeared stronger than that exerted by Dkstatin-2 (Fig 7A). Again, among those genes whose expression was substantially altered, only a small portion was simultaneously affected by either compound.

**Fig 7.**
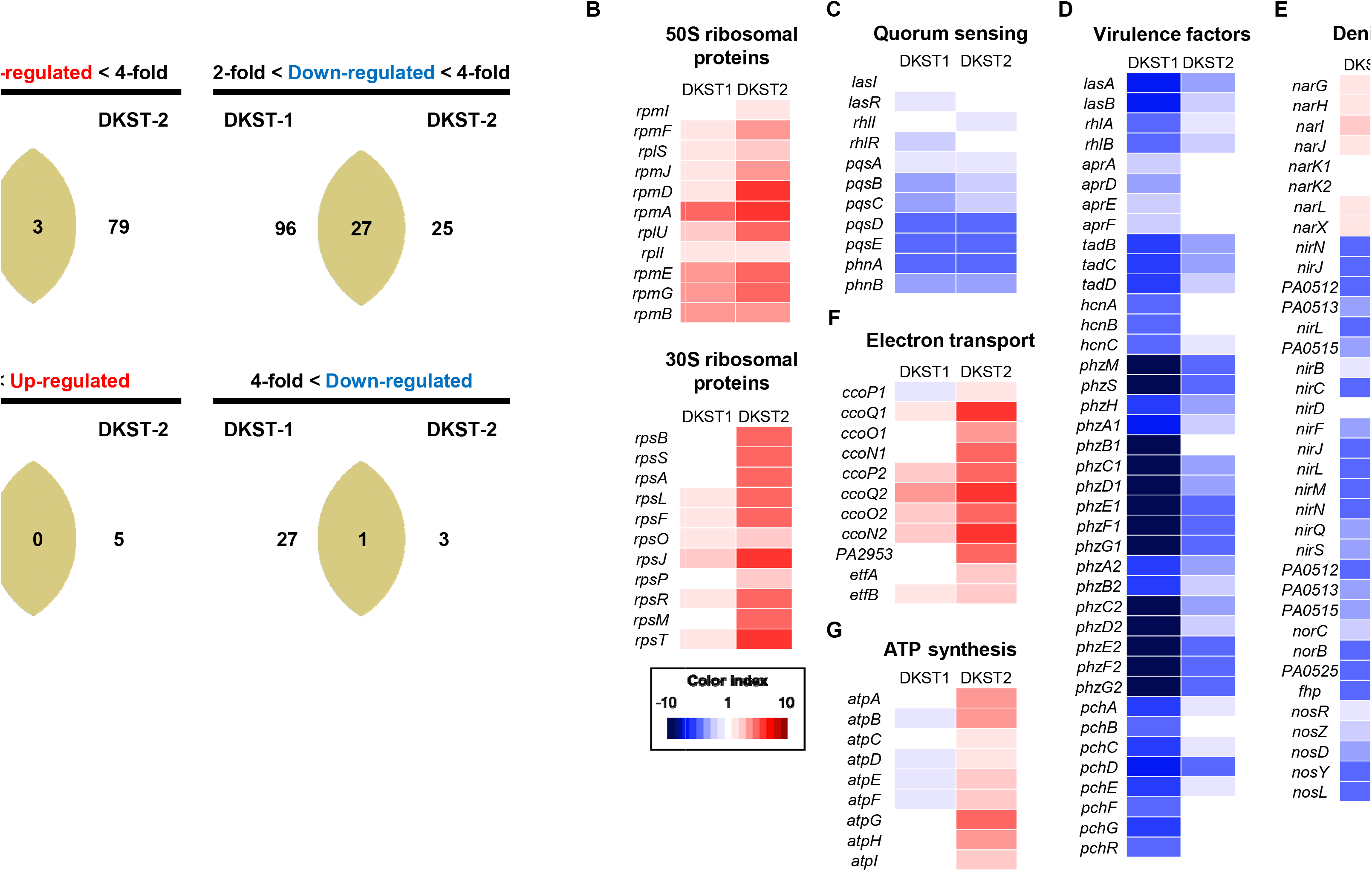
Comparison of effects of Dkstatin-1 and Dkstatin-2 on gene expression. (**A**) Venn diagram for number of genes affected by Dkstatin-1 (DKST-1) or Dkstatin-2 (DKST-2) treatment. Expression fold range was separated as 2 to 4-fold and over 4-fold. (**B**) Expression of 50S and 30S ribosomal protein encoding genes. Dkstatin-2 (DKST-2) was more effective to increase expression of those genes. (**C**) Expression of genes participated in three major QS circuits was downregulated in the presence of both Dkstatin-1 and Dkstatin-2. (**D**) Transcription of genes for virulence factors. Phenazine biosynthesis clusters (*phzA1~G1* and *phzA2~G2*) were remarkably down-regulated for either Dkstatin. (**E**) Expression of genes involved in all steps of denitrification. Gene clusters (*nirSMCFLGHJND* and *norCBD*) encoding nitrite reductase (NIR) and nitric oxide reductase (NOR) were invariably down-regulated in both Dkstatin-1 and Dkstatin-2. (**F**) Expression of genes categorized as electro transfer. The two cbb3-type cytochrome c oxidase encoding gene set (*ccoN1~P1* and *ccoN2~P2*) was up-regulated by Dkstatin-2. (**G**) Expression of ATP synthase complex encoding gene operon. All total RNA were harvested as biological duplicates.

At first, we confirmed that the transcription of genes encoding ribosomal proteins was upregulated during the treatment with Dkstatin-1 or Dkstatin-2, with the latter exhibiting more significant effects (Fig 7B). These results further verified that two compounds can in fact effectively compromise the activity of DksA1. We then examined the effects of Dkstatin treatment on the expression of genes involved in QS and QS-mediated virulence factor production. As shown in Fig 7C, transcription of genes encoding autoinducer synthases and autoinducer-binding receptors was suppressed in Dkstatin-treated PAO1 cells. Among these, *pqsABCDE* and *phnAB* genes that constitute the PQS biosynthesis cluster were most affected. Importantly, the expression of virulence-associated genes was remarkably decreased, especially in response to the treatment with Dkstatin-1 (Fig 7D). Genes coding for elastase (*lasB*), alkaline protease (*aprADEF*) and rhamnolipid (*rhlAB*) were considerably downregulated. Similarly, transcription of genes involved in phenazines production (*phzA1~G1* and *phzA2~G2*) and siderophore biosynthesis (*pchA~G* and *pchR*) were also suppressed in Dkstatin-treated cells (Fig 7D). These results clearly suggest that Dkstatin-induced virulence suppression is due to the effects manifested at the transcriptional level. In addition, gene expression for the production of hydrogen cyanide (*hcnA~C*), and type II secretion machinery (*tadB~D*) was less active in the presence of Dkstatins.

In our previous data shown in Fig 4A, Dkstatin treatments suppressed anaerobic respiratory growth of PAO1. In *P. aeruginosa*, anaerobic respiration is mediated by a series of reductions from nitrate (NO_3_^-^) to N_2_ with nitrite (NO_2_^-^), nitric oxide (NO) and nitrous oxide (N_2_O) as intermediates [25]. The expression of genes involved in the activity of nitrate reductase (NAR) that mediates the first step of anaerobic denitrification was slightly increased in the presence of Dkstatins. In contrast, Dkstatins significantly suppressed the expressions of gene clusters (*nirSMCFLGHJND* and *norCBD*) encoding nitrite reductase (NIR) and nitric oxide reductase (NOR) that catalyze the next steps of denitrification to produce nitrous oxide [26]. Similarly, the expression of *nosRZDFYL* cluster encoding nitrous oxide reductase also invariably decreased during Dkstatin treatment (Fig 7E).

Our data shown in Fig 2A indicate that Dkstatin-2 stimulated PAO1 growth in LB. To address this unexpected phenomenon, our interest also reached the genes involved in energy metabolism such as systems for electron transport and ATP biosynthesis. The two gene clusters, *ccoN1O1Q1P1* and *ccoN2O2Q2P2*, encode subunits of cbb3-type cytochrome oxidase, a critical enzyme for aerobic respiration [27, 28]. Compared to the control PAO1 cells, transcription of these two gene clusters was remarkably increased in PAO1 cells treated with Dkstatin-2 (Fig 7F). Furthermore, a large operon *atpABCDEFGHI* encoding ATP synthase complex was more actively transcribed in the same cells (Fig 7G). These results provided insight into how Dkstatin-2 treatment promoted the PAO1 aerobic growth. Collectively, our RNASeq analysis revealed that two Dkstatin compounds induced transcriptional changes, which were well reflected at the phenotypic level. In addition, we further confirmed that the wild type PAO1 cells subjected to Dkstatin treatment behaved like a Δ*dksA1* mutant.

### Dkstatin-1 and Dkstatin-2 attenuated virulence of *P. aeruginosa in vivo*

As the production of virulence factors decreased by Dkstatin treatment, we next tested whether Dkstatin-induced virulence attenuation is still valid *in vivo* using 8-week-old C57BL/6 female mice. All mice were intranasally infected with freshly harvested PAO1 cells (~10^6^ CFU). Mice infected with PAO1 (n=3) perished within 36 hours (Fig 8A, green line). On the other hand, when either Dkstatin was supplemented together with PAO1 infection (n=4), mice exhibited significantly increased survival rates and survived even after at 48 hour post-infection (Fig 8A). In a separate set of experiments, we also examined the effect of Dkstatin treatment on PAO1 proliferation inside the host airway. In this case, mice were infected with 3×10^6^ PAO1 cells. At 12 hour post-infection, the average bacterial load in the lungs of the control group was ~5.2×10^5^ CFU. Importantly, the bacterial load was significantly decreased to ~4.7×10^4^ CFU, when treated with Dkstatin-1. In contrast, the bacterial count was not decreased by Dkstatin-2 (Fig 8B). This finding correlates well with our RNASeq results, in that Dkstatin-1 treatment strongly suppressed the transcription of virulence-associated genes (Fig 7F). We also assessed the effects of Dkstatin treatment by visualizing lung tissues. Intranasal infection of PAO1 led to infiltration of immune cells and a reduction of alveolar spaces, which are indicators of severe airway infections (Fig 8F). When PAO1-infected mice were treated with Dkstatins, inflammatory symptoms substantially decreased compared with those observed in the control infection group (Fig 8G and H vs. Fig 8F). The presence of lung damage was not obvious when the mice were treated with Dkstatin-1 (Fig 8D) or Dkstatin-2 (Fig 8E). Together, these results suggested that Dkstatin-1 and Dkstatin-2 are not toxic to the murine host and thereby these two molecules may have potential to be developed as anti-Pseudomonas agents for human patients.

**Fig 8.**
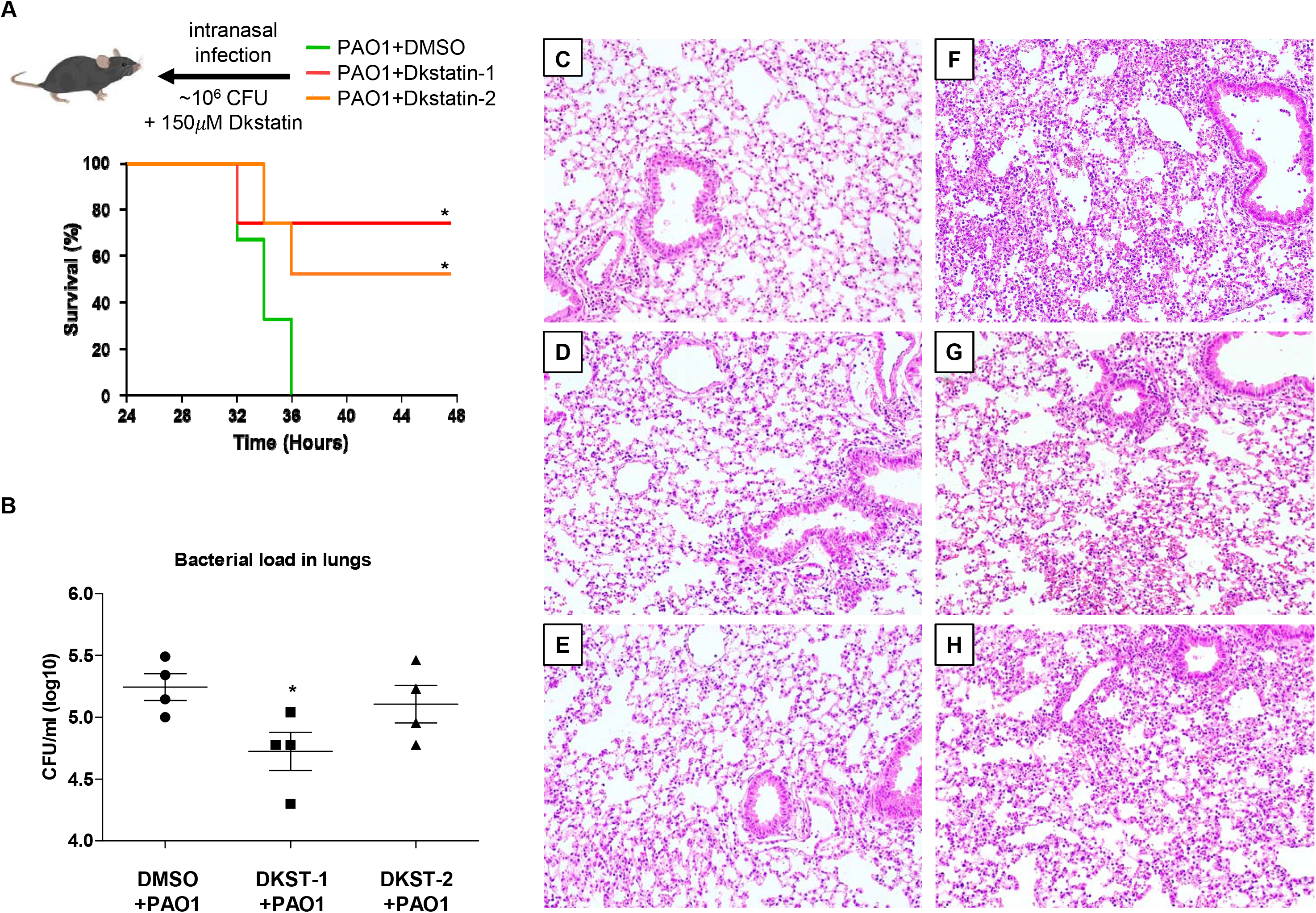
*in vivo* virulence analyses of Dkstatin-1 and Dkstatin-2. (**A**) Survival curves of mice infected with PAO1 (green line) and PAO1 combined with Dkstatin-1 (red line) or Dkstatin-2 (orange line). The infection dose was ~10^6^ cells per mouse. *, *p* < 0.05 *vs*. survival rate of PAO1-infected mice. Four mice were used in each group. (**B**) Bacterial CFU values counted from whole dissected lungs. The initial dose was 3 x 10^6^ cells per mouse of PAO1 or PAO1 combined with 150 μM of Dkstatin-1 or Dkstatin-2. *, *p* < 0.05 *vs*. CFU of the PAO1-infected mice. (**C**) H&E stained histological visualizations of lung tissues from mice intranasally administrated DMSO only, (**D**) 150 μM of Dkstatin-1 and (**E**) Dkstatin-2. (**F**) H&E stained lung tissues of mice infected with PAO1 combined with DMSO, (**G**) PAO1 combined with 150 μM of Dkstatin-1 and (**H**) Dkstatin-2.

## Discussion

Previous studies from our lab revealed that DksA1 acts as a major transcriptional regulator for QS-mediated virulence expression in *P. aeruginosa* [13]. In addition, DksA1 is also required for uninterrupted expression of genes encoding proteins for biofilm formation, motility, anaerobic respiration through denitrification and polyamine metabolism [13]. As DksA1 participates in regulating a wide range of phenotypes, including virulence in *P. aeruginosa*, we postulated that DksA1 is an effective drug target, the inhibition of which may result in virulence attenuation, eventually leading to control of recalcitrant *P. aeruginosa* infections. The success of our compound library screen is attributed to the following aspects. First, the use of formazan production as a surrogate marker of DksA1activity provided a system that allowed us to screen 6,970 compounds with minimal error. Second, the reporter strain, in which the transcriptional fusion of *P_rpsB_::lacZ* has been inserted as a single copy at a nonfunctional region of PAO1 genome [29], provided a reliable method for the second-stage verification. This was critical because reduced production of formazan can result from other metabolic alterations. Lastly, Dkstatin-1 and Dkstatin-2 were found to be commercially available for purchase, enabling us to test the effects of these two final candidates at higher concentrations with no limited supply. Considering that the initial screening could only be performed with compounds at 50 μM concentration, the availability of Dkstatins made this study more feasible.

Dkstain-1 and Dkstain-2 have quite different chemical structures. Both compounds, however, induced specific outcomes mediated by loss of DksA1 activity, ranging from (i) increased *rpsB* expression, (ii) reduced elastase production, (iii) reduced pyocyanin production, (iv) suppressed autoinducer production and (v) defective anaerobic respiration. These results display the common features of two compounds affecting DksA1 activity. DksA1 possesses a Zn^2+^ binding motif (CXXC-N^17^-CXXC) consisting of 4 cysteine residues at its C-terminus, and this motif is critical for its structure and function. Of note, the cysteine thiol group is highly susceptible to oxidation by electrophilic molecules [3, 19, 30]. Regardless of the structural differences, both compounds harbor reactive groups, such as fluorine [31] and butanone [32, 33]. Thus, it is reasonable to hypothesize that oxidizing groups in Dkstatins may affect the DksA1 structure, which in turn leads to interference with DksA1 binding to a subunit of RNAP. Future work should focus on understanding Dkstatin-induced structural modification of DksA1 and its downstream consequences. To this end, it is necessary to explore other chemicals harboring diverse oxidizing moieties as well.

Based on Fig 2A, Dkstain-2 stimulated bacterial growth. While the enhanced bacterial growth was not fully reflected in the viable cell counting assay, we became interested in this unexpected result. It is counterintuitive that a compound that has potential to be developed as an anti-infective agent also stimulates bacterial growth. However, it is also noteworthy that actively replicating cells are well known to be more susceptible to antibiotic treatment [34]. In line with this notion, *P. aeruginosa* invaders might be more exposed to host immunity when they multiply at a faster rate. In contrast to the aerobic culture, Dkstatin-2 strongly suppressed anaerobic respiratory growth of PAO1 (Fig 4B). Earlier studies clearly suggest that *P. aeruginosa* may benefit from anaerobiosis during chronic airway infection [25]. It will be important to further clarify whether Dkstatin-2 specifically suppresses chronic airway infection by *P. aeruginosa*.

QS, a cell density-dependent gene regulatory system, is crucial for the production of virulence determinants in *P. aeruginosa* [22]. Therefore, *P. aeruginosa* QS has been extensively studied as a therapeutic target for virulence attenuation [1, 22, 35]. To date, strategies for QS inhibition include receptor inactivation [36], inhibition of signal synthesis, use of autoinducer analogues [37], signal degradation [38, 39] and use in combination with antibiotics [37]. Flavonoids bind QS receptor, LasR and RhlR, to reduce virulence factor production [36]. N-decanoyl-L-homoserine benzyl ester, a structural analogue of 3-oxo-C12-HSL, downregulates elastase production, swarming motility and biofilm formation without growth defects [21]. This QS-inhibitor also synergistically interacts with gentamicin and tobramycin [37]. Another QS inhibitor, meta-bromo-thiolactone, affects LasR and RhlR activity to decrease pyocyanin production and biofilm formation in *P. aeruginosa* PA14 strain [40]. In a screen with a synthetic N-acyl-homoserine lactone (A-HSL) library, various HSL derivatives were found to antagonize the activity of LasR and LuxR in *P. aeruginosa* and *Vibrio fischeri* [41]. Likewise, a couple of autoinducer analogues were found to antagonize TraR, a QS-related LuxR homologue in *Agrobacterium tumefaciens* [41]. Of note, the previously mentioned examples show findings from studies designed to uncover molecules that directly inhibit or antagonize at various levels of the *P. aeruginosa* QS machinery. In the present study, however, we discovered molecules that target DksA1, whose involvement in *P. aeruginosa* QS is relatively a new subject. We argue that DksA1 should be an avenue for future exploration in the context of development of *P. aeruginosa* QS inhibitors with a focus on its molecular nature and its impact on virulence gene regulation.

Our RNASeq analysis clearly suggested that the two Dkstatins induced distinct changes in the genome-wide transcription pattern. When differentially expressed genes (DEGs) were categorized based on the gene ontology (GO) term, a larger number of GO terms were retrieved in Dkstatin-2 treated group (11 terms) than in the Dkstatin-1 treated group (9 terms). Of particular interest, the most significantly presented GO term in the Dkstatin-2 treated group was “translation” with all 23 upregulated genes (S5B Fig). Along with this GO term, several other functional categories, such as ribosomal assembly and energy metabolism, were also identified with genes of enhanced transcription. Given that these changes resemble the characteristics of the Δ*dksA1* mutant, it is likely that Dkstatin-2, as compared with Dkstatin-1, is more potent in inhibiting DksA1 activity. Consistent with this idea, binding of DksA1::FLAG and RpoB was hampered to a greater degree by Dkstatin-2 (Fig 6B). In the Dkstatin-1 treated group, on the other hand, the “phenazines biosynthesis process” was most significantly presented (S5A Fig). In addition, 10 out of 12 genes in the “pathogenesis” category and all genes in the “secretion”, “type VI secretion system” and “chemotaxis” terms were downregulated in Dkstatin-1 treated cells (S5A Fig), suggesting that Dkstatin-1 may be more directly engaged in attenuating *P. aeruginosa* virulence. Collectively, these additional bioinformatic analyses further support the notion that Dkstatin-1 and Dkstatin-2 may operate in different modes of action even though these two molecules were identified as DksA1 inhibitors.

Antibiotic resistance in many human pathogens (including *P. aeruginosa*) has already become a global healthcare problem. Against the opportunistic features of *P. aeruginosa*, we need to come up with antibiotic-independent infection control strategies. Interventions that could achieve virulence attenuation without imposing selective pressure have been actively attempted [35]. Dkstatins identified in the present study significantly downregulated *P. aeruginosa* virulence by suppressing QS mechanisms. Moreover, Dkstatins induced elevated antibiotic efficacy at sub-MIC conditions, especially for antibiotics that inactivate protein synthesis. As DksA1 suppresses transcription of genes encoding rRNA and ribosomal proteins, macromolecular ribosomes are readily accumulated when DksA1 is inactivated. Therefore, it is anticipated that ribosome-targeting antibiotics are more active under such conditions.

In our animal model of acute infection, two Dkstatin molecules successfully protected mice from a lethal dose of PAO1 infection. Provided that Dkstatins are safe in humans, they will broaden our options to treat *P. aeruginosa* infections with low risk of antibiotic resistance. Collectively, Dkstatins are attractive drug candidates in establishing future plans to cope with antibiotic-resistant infections. We anticipate that future investigations will propose detailed molecular mechanisms of Dkstatins and apply these in human trials.

## Methods

### Ethics statement

All animal studies were performed in compliance with the guidelines provided by the Department of Animal Resources of Yonsei Biomedical Research Institute. The Committee on the Ethics of Animal Experiments at the Yonsei University College of Medicine approved this study (permit number 2018-0246).

### Bacterial strains, culture conditions and chemicals

Bacterial species and plasmids used in this study are listed in S1 Table. A laboratory strain of *P. aeruginosa*, PAO1, was used as a wild type strain. Unless otherwise specified, bacterial strains were cultivated in Luria-Bertani media (LB; 1% (w/v) tryptone, 0.5% (w/v) yeast extract, and 1% (w/v) sodium chloride) at 37°C. Standard cloning procedures that involved allelic exchange were performed. When necessary, ampicillin (100 μg/ml), carbenicillin (100 μg/ml), gentamycin (60 μg/ml), or irgasan (25 μg/ml) were used for clonal selection. DNA sequences encoding 3XFLAG tag were amplified and ligated to the downstream of the multi-cloning site (MCS) of the pBAD24 plasmid, resulting in the construction of pBAD24F. Then, the *dksA1* gene was cloned into the MCS of pBAD24F. The resulting *dksA1::FLAG* sequence was amplified and ligated into pCVD442 [34] for chromosomal integration. The promoter region of *rpsB* gene was fused with the promoter-less *lacZ* open reading frame (P_*rpsB*_::*lacZ*), and the resulting construct was cloned into the plasmid, pUC18-mini-Tn7t-Gm-*lacZ* [29] for chromosomal integration. Dkstatin-1 (4,4’-(4,4’-biphenyldiyl disulfonyl)di(2-butanone)) and Dkstatin-2 (N-ethyl-3-[(3-fluorophenyl)methoxy]-N-[(1-methyl-1H-imidazol-5-yl)methyl]aniline) were purchased from Chembridge Corp. (San Diego, CA) and Asinex Inc. (Amsterdam, Netherlands), respectively.

### Screening of Chemical Compounds

Screening of chemical compounds was performed in two phases. Using a chemical compound library consisting of 6,970 chemicals provided by the Korea Chemical Bank (https://chembank.org/), we first screened for chemical compounds that inhibit DksA1 activity. The total library was 10-fold diluted with DMSO to increase the volume. A 100-fold diluted overnight culture of *P. aeruginosa* PAO1 and the Δ*dksA1* mutant were then incubated in fresh media for an additional 4 hours to activate the cells. The obtained OD_600_ values of these strains were adjusted to 1.0. Then, 120 μl of the adjusted culture was inoculated in 96-well plates that contain 50 μM of chemical compounds, and these were incubated at 37°C for 1 hour to examine formazan production. After incubation, 0.05 mg/ml of thiazolyl blue tetrazolium bromide (Sigma-Aldrich) was distributed into each well of the test plates, and they were incubated for an additional 30 min at 37°C. The formazan production was measured at a 550 nm wavelength after dissolving it in DMSO. The top 50 compounds that presented the lowest levels of formazan production during the first screening phase were then subjected to a subsequent screening. In the following screening phase, a reporter strain PAO1 *P_rpsB_::lacZ* was diluted 100-fold and added to 1.5 ml of LB supplemented with 50 μM of each of the 50 compounds. The sample was then incubated for 4 hours at 37°C with shaking. *rpsB* expression of the reporter strain was represented in Miller Units and was measured as described previously [13]. Based on the *rpsB* expression, the top 20 compounds among the 50 previously screened compounds were selected. To further narrow down the candidate compounds from the 20 selected compounds, the reporter strain was incubated in 3 ml of LB with the addition of 50 μM of each of the 20 compounds for 4 hours with shaking. Measurement of the elastase activity and quantification of *rpsB* expression allowed us to come down to the 4 most efficacious compounds: 55B05, 02G09, 86B09, and 45G08. Then, 50 μM of four these compounds were tested again by measuring elastase activity and *rpsB* expression of reporter strain incubated in 5 ml of LB for 4 to 8 hours at 37°C. 55B06 and 86B09 were specifically chosen as they showed sustained effects on elastase activity and *rpsB* expression. Then, 150 μM of 55B05 and 86B09 were each added to PAO1 and Δ*dksA1* mutant incubated in 5 ml of LB medium for 8 hours at 37°C to measure formazan production.

### Elastase and pyocyanin assay

Elastase activity was measured as previously described [42]. First, 500 μl of supernatant from cultures incubated with 50 μM or 150 μM of the compounds for 4 hours or 8 hours was mixed with 1 ml of 30 mM Tris-HCl buffer containing 10 mg/ml of Elastin-Congo Red (Sigma-Aldrich). The mixture was then incubated with shaking at 37°C for 1–2 hours. After the incubation, 1 ml of the mixture was transferred into a micro tube, which was centrifuged at 13,000 rpm for 1 min. In the pyocyanin assay, 5 ml of supernatants from the cultures incubated for 8 hours with 150 μM Dkstatin compounds were harvested by centrifugation at 3,000 rpm for 20 min. They were then filter-sterilized (0.2-μm pore size, Sartorius Minisart Sterile EO filters; Sartorius AG). Then, 4 ml of chloroform was added to 8 ml of the supernatants in a 15 ml conical tube, and the mixture was homogenized. After homogenization, the tubes were centrifuged at 3,000 rpm for 10 min to collect the blue layer fraction submerged to the bottom of the tube. Then, 4 ml of the collected blue layer was transferred into a new tube, and 2 ml of 0.2 N HCl was added to produce a red colored mixture. For the assay, 1 ml of the mixture was then transferred into a microtube for centrifugation at 13,000 rpm for 1 min. Absorbance at 520 nm (OD_520_) was then measured for the samples.

### Autoinducer assay

In HSL measurements, both C4-HSL and 3-oxo-C12-HSL were extracted from 5 ml of culture supernatants using the equivalent volume of acidified ethyl acetate. The extractions were repeated twice to fully collect the ethyl acetate fraction. N_2_ gas was used to evaporate and dry out the collected ethyl acetate fraction. The residues were dissolved in 250 μl of HPLC grade ethyl acetate to obtain 20-fold concentrated extracts. Measurement of 3-oxo-C12-HSL was performed using an *E. coli* strain harboring reporter plasmid pKDT17 containing a copy of *lasR-lacZ* transcriptional fusion [43]. Samples were 100-fold diluted to culture a volume of reporter strain, and they were then tested to measure β-galactosidase activity. The Miller Units from samples were calculated as described previously [13]. The measurement of C4-HSL was conducted through violacein productions of the *Chromobacterium violaceum* CV026 strain [44]. The CV026 strain was incubated in 5 ml of LB and LB supplemented with 100-fold diluted extract. After incubation, cell pellets in 1 ml of culture were harvested by centrifugation at 13,000 rpm for 3 min. The pellets were then re-suspended using HPLC grade DMSO to dissolve violacein. Violacein was measured at 575 nm absorbance and normalized using the OD_600_ value of CV026.

### DNA manipulation

Primers used in the study are listed in S2 Table. Plasmid preparation was performed using a Plasmid mini extraction kit (Bioneer Inc., Seoul). Restriction enzyme digestion, ligation, and agarose gel electrophoresis were performed by following standard methods. A Gibson assembly kit (New England Biolabs Inc., MA.) was used in cloning to produce the *dksA1*-fused 3XFLAG strain. Transformation of *E. coli* was carried out by electroporation. Competent *E. coli* cells for electroporation were prepared by repeated washing with 300 mM sucrose. Electroporation settings were 2.5kV, 25μF and 200 Ω for a 2 mm electroporation cuvette. The oligonucleotide primers synthesis and sequencing DNA were performed by Macrogen, Korea.

### Antibiotic susceptibility test

Overnight cultured PAO1 was inoculated to 100-fold dilution in 2 ml of LB supplemented with gentamicin, imipenem, streptomycin, tetracycline, kanamycin and tobramycin ranging from 1 μg/ml to 25 μg/ml, and 150 μM of Dkstatin-1 or Dkstatin-2. The incubation took 6 hours at 37°C with vigorous shaking. Then, respective incubated samples were serially diluted by 10-fold in 1 ml of PBS, and 10 μl of each dilute was spotted onto an LB agar plate for CFU counts. The plates were incubated overnight at 37°C without shaking.

### Western blot and immunoprecipitation

PAO1 *dksA1*::3xFLAG strain was grown in 20 ml of LB medium supplemented with or without 150 μM of Dkstatin-1 and Dkstatin-2 for 6 to 8 hours. Whole cells were pelleted by centrifugation at 3,000 rpm for 30 min at 4°C and were re-suspended with 600 to 800 μl of lysis buffer (50 mM Tris-HCl pH 7.5, 75 mM NaCl, 10 mM MgCl2, 10% glycerol, 1 mM EDTA, 1 mM PMSF and 0.01% Triton X-100) with 8 μl of SIGMA FAST protease inhibitor cocktail (Sigma-Aldrich). The pellets were then sonicated at 30% amplitude for 10 sec with 10 sec intervals on ice and were then centrifuged for 30 min at 4°C and 14,000rpm. The supernatants of the lysates were collected and placed into new tubes. Their volumes were adjusted to contain equal quantities of total proteins. Then, 10 to 800 μl of the adjusted samples were used for western blots and immunoprecipitation. For DksA1::3xFLAG blotting, 10 μl of samples were loaded on 12% SDS-PAGE and transferred to Amersham Protran premium 0.2 μm nitrocellulose membranes (GE Healthcare Life Sciences) for 1 hour in an ice-cold transfer buffer (25mM Tris, 192mM glycine pH 8.3 20% methanol (v/v)). Membranes were blocked with a 5% BSA containing TBST solution overnight at 4°C on a rocking shaker. Blots were incubated with anti-FLAG (Sigma-Aldrich) and anti-rabbit secondary antibodies (Thermo-fisher) and revealed by chemi-luminescence. All washes were performed with a TBST solution without BSA. Immunoprecipitation was conducted using ~800 μl of samples with 25 μl of M2 Anti-FLAG magnetic beads (Sigma-Aldrich) after incubating overnight at 4°C on a rocking shaker. Beads were pelleted using a magnet and were washed once with the lysis buffer. Samples bound to the beads were eluted in 30 μl of lysis buffer by heating at 100°C for 10 min. The eluted samples were then loaded on 10% SDS-PAGE and blotted by anti-FLAG Ab (Sigma) or anti-*E. coli* RpoB Ab (Abcam).

### RNASeq analysis

To extract total RNA, *P. aeruginosa* PAO1 was incubated for 2 hours in 5 ml of LB with vigorous shaking at 37°C. Then, additional incubation was conducted for 1 hour under the same condition with 150 μM of Dkstatin-1 or Dkstatin-2 supplement. Total RNA samples were extracted using an RNeasy mini kit (Qiagen), and total DNA was eliminated using RNase-free DNase set (Qiagen) following the manufacturer’s instructions. The extraction was performed as an experimental duplicate. Transcriptome profile comparison and processing the expression data were provided by Ebiogen (Korea). RNA quality was assessed by an Agilent 2100 bioanalyzer using the RNA 6000 Nano Chip (Agilent Technologies, Amstelveen, The Netherlands), and RNA quantification was performed using an ND-2000 Spectrophotometer (Thermo Inc., DE, USA). For control and testing RNAs, rRNA was removed using a Ribo-Zero Magnetic kit (Epicentre Inc., USA) from each 5 ug of total RNA. The construction of a library was performed using a SMARTer Stranded RNA-Seq Kit (Clontech Lab Inc., CA, USA) according to the manufacturer’s instructions. The rRNA depleted RNAs were used for the cDNA synthesis and shearing, following manufacturer’s instructions. Indexing was performed using the Illumina indexes 1–12. The enrichment step was carried out using PCR. Subsequently, libraries were checked using the Agilent 2100 bioanalyzer (DNA High Sensitivity Kit) to evaluate the mean fragment size. Quantification was performed using the library quantification kit using a StepOne Real-Time PCR System (Life Technologies Inc., USA). High throughput sequencing was performed as paired-end 100 sequencing using HiSeq 2500 (Illumina Inc., USA). Bacterial-Seq reads were mapped using a Bowtie2 software tool to obtain the alignment file. Differentially expressed genes were determined based on counts from unique and multiple alignments using EdgeR within R (R development Core Team, 2016) using Bioconductor [45]. The alignment file also was used for assembling transcripts. The quantile normalization method was used for comparisons between samples. Gene classification was based on searches done by DAVID (http://david.abcc.ncifcrf.gov/).

### Mouse infection

Eight-week-old female C57BL/6 mice (Orient, Korea) were infected with PAO1 and a combination of Dksatin-1 or Dkstatin-2. PAO1 pellets incubated in LB broth at 37°C for 6 hours were resuspended in 1 ml of sterile phosphate-buffered saline (PBS) containing 1% of DMSO or 150 μM of Dkstatin compounds, and they were adjusted to 10^6^ cells per 20 μl as an initial infection dose. Mice were anesthetized by injection of 20% (v/v) ketamine and 8% (v/v) Rompun mixed in saline. During anesthetization, 20 μl of prepared bacterial cells were inhaled directly through the nose by a pipette as previously described [46]. The survival of the mice was monitored for 48 hours with 2 hour intervals. The survival of the mice was expressed as a Kaplan-Meier curve using Graph Pad Prism software (www.graphpad.com). To measure the bacterial load in the lungs, another set of mice (n=4) were infected with the same administration above. All mice were euthanized in a CO_2_ gas chamber 12 hours post infection. Then, whole lungs from mice in each group were immediately collected and homogenized for serial dilution in 1 ml of PBS. For histological observation, lungs were perfused with sterile PBS and staining with hematoxyline and eosin was conducted following a standard protocol [47].

### Statistical analysis

Statistical analysis of the data in the experiments was carried out by statistic tools within Graph Pad Prism software (www.graphpad.com) for the paired Student’s *t*-test. A *p*-value of less than 0.05 was considered statistically significant. All experiments were repeated for reproducibility.

## Acknowledgements

This work was supported by grants from the National Research Foundation (NRF) of Korea, which is funded by the Korean Government (2017M3A9F3041233 and 2019R1A6A1A03032869). This research was also supported by a grant from the Korea Health Technology R&D Project through the Korea Health Industry Development Institute (KHIDI), funded by the Ministry of Health & Welfare, Republic of Korea (HI14C1324). We would like to thank Jun Bum Kim and Jiwon Kim for proofreading the manuscript. The chemical library used in this study was kindly provided by Korea Chemical Bank (www.chembank.org) of Korea Research Institute of Chemical Technology.

## Author Contributions

K.B.M., W.H., K.M.L. and S.S.Y. conceptualized and designed the experiments. K.B.M and S.S.Y performed experiments and analyzed the experimental results. K.B.M. and S.S.Y drafted the manuscript.

## Supplementary figure legends

**S1 Fig. Screening scheme for Dkstatin compound. (A)** Visual comparison of formazan production. Reduction of formazan production in Δ*dksA1* mutant was observed. **(B)** Demonstration of screening procedure for Dkstatin compound. In the first step, the OD600 value of *P. aeruginosa* incubated in LB for 4 hours was adjusted to 1.0, and it was inoculated into chemical library plates. 15 μl of chemical compounds in stock chemical libraries were distributed into new plates to at a concentration of 50 μM. Exposure of chemical compound to PAO1 was performed for 2 hours at 37°C. 15 μl of 0.5 mg/ml thiazolyl blue tetrazolium bromide was inoculated into test plates to observe (purple colored) formazan production. In the second step in the screening, *rpsB* expression (represented as β-galactosidase activity) and elastase activity were measured to select further potent chemical candidates.

**S2 Fig. Measurements of relative elastase activity and *rpsB* expression under four Dkstatin candidates (A)** Comparisons of relative elastase activity and *rpsB* expression from PAO1 *PrpsB::lacZ* and PAO1Δ*dksA1 PrpsB::lacZ* strains incubated for 4 hours in LB. Relative levels of *rpsB* expression were represented as β-galactosidase activity. 50 μM of 4 Dkstatin candidate compounds (labeled as 55B05, 02G09, 86B09 and 45G08) were supplemented in LB. The values of mean ± S.D. are presented (n=2). **(B)** Relative levels of elastase activity and *rpsB* expression of *PrpsB::lacZ* and PAO1Δ*dksA1 PrpsB::lacZ* strains incubated for 8 hours in LB. Relative levels of *rpsB* expression were represented as β-galactosidase activity. 50 μM of 4 Dkstatin candidate compounds were supplemented in LB. The values of mean ± S.D. are presented (n=2).

**S3 Fig. HSL-autoinducer complementation under Dkstatin-1 and Dkstatin-2 supplemented condition.** Demonstration of combined HSL-autoinducer (C4/C12-HSL) complementation experiment procedure (Upper). 200 μM of C4/C12-HSL complementation in PAO1 and Δ*dksA1* mutant was conducted for 8 hours in LB medium with supplementation of 150 μM of Dkstatin-1 or Dkstatin-2. Elastase activities of PAO1 and Δ*dksA1* mutant were measured at each time point after addition of C4/C12-HSL (4, 6, and 8 hours). Elastase production in PAO1 treated with 150 μM of Dkstain-1 (o, blue line) and Dkstatin-2 (o, orange line) was constantly less than 50% that produced in the control (•, black line). With C4/C12-HSL complementation, the elastase productions with 150 μM of Dkstain-1 (•, red line) and Dkstatin-2 (•, green line) were restored in 4 hours and completely diminished 8 hours post complementation. C4/C12-HSL complementation was not effective in elastase production in Δ*dksA1* mutant (□, black line and ■, black line).

**S4 Fig. Transcriptome analysis produced in PAO1 and PAO1 treated with Dkstatin-1 or Dkstatin-2.** (**A**) Transcriptome heat-map produced from biologically duplicated RPKM values of PAO1 and Dkstatin-1 or −2 treated PAO1. DMSO was used as a solvent control for Dkstatin-1 and Dkstatin-2 treatment. The Z-score ranged from −1.0 to 1.0 and is represented as a color coded index. (**B**) PCA plot representing the similarity of transcripts in PAO1 treated with DMSO (#1 and #2) and with Dkstatin-1 (#5 and #6) or Dkstatin-2 (#3 and #4). Each similarity was significantly distinguishable. Total transcripts were harvested as biological duplicates.

**S5 Fig. Categorization of differentially expressed genes based on gene ontology (GO) terms.** (**A**) GO terms in Dkstatin-1 treated group. Most genes included in GO terms (such as phenazines biosynthesis process, pathogenesis, secretion and chemotaxis) were down-regulated by Dkstatin-1 treatment. (**B**) Presented GO terms in Dkstatin-2 treated group. 11 GO terms were retrieved in the Dkstatin-2 treated group. Genes participating in GO terms such as translation, ribosomal assembly and energy metabolism were up-regulated by Dkstatin-2 treatment.

**S1 Table. Bacterial strains and genetic materials used in this study.**

**S2 Table. Primer sequences used in this work.**

## References

1. Moradali MF, Ghods S, Rehm BH. Pseudomonas aeruginosa Lifestyle: A Paradigm for Adaptation, Survival, and Persistence. Front Cell Infect Microbiol. 2017;7: 39. doi:10.3389/fcimb.2017.00039 PMID:28261568

2. Valentini M, Gonzalez D, Mavridou DA, Filloux A. Lifestyle transitions and adaptive pathogenesis of Pseudomonas aeruginosa. Curr Opin Microbiol. 2018;41: 15–20. doi:10.1016/j.mib.2017.11.006 PMID:29166621

3. Gourse RL, Chen AY, Gopalkrishnan S, Sanchez-Vazquez P, Myers A, Ross W. Transcriptional Responses to ppGpp and DksA. Annu Rev Microbiol. 2018;72: 163–184. doi:10.1146/annurev-micro-090817-062444 PMID:30200857

4. Ross W, Sanchez-Vazquez P, Chen AY, Lee JH, Burgos HL, Gourse RL. ppGpp Binding to a Site at the RNAP-DksA Interface Accounts for Its Dramatic Effects on Transcription Initiation during the Stringent Response. Mol Cell. 2016;62: 811–823. doi:10.1016/j.molcel.2016.04.029 PMID:27237053

5. Doniselli N, Rodriguez-Aliaga P, Amidani D, Bardales JA, Bustamante C, Guerra DG, et al. New insights into the regulatory mechanisms of ppGpp and DksA on Escherichia coli RNA polymerase-promoter complex. Nucleic Acids Res. 2015;43: 5249–5262. doi:10.1093/nar/gkv391 PMID:25916853

6. Paul BJ, Barker MM, Ross W, Schneider DA, Webb C, Foster JW, et al. DksA: a critical component of the transcription initiation machinery that potentiates the regulation of rRNA promoters by ppGpp and the initiating NTP Cell. 2004; 118: 311–322. doi:10.1016/j.cell.2004.07.009 PMID:15294157

7. Perederina A, Svetlov V, Vassylyeva MN, Tahirov TH, Yokoyama S, Artsimovitch I, et al. Regulation through the secondary channel--structural framework for ppGpp-DksA synergism during transcription. Cell. 2004;118: 297–309. doi:10.1016/j.cell.2004.06.030 PMID:15294156

8. Bernardo LM, Johansson LU, Solera D, Skarfstad E, Shingler V. The guanosine tetraphosphate (ppGpp) alarmone, DksA and promoter affinity for RNA polymerase in regulation of sigma-dependent transcription. Mol Microbiol. 2006;60: 749–764. doi:10.1111/j.1365-2958.2006.05129.x PMID:16629675

9. Kolmsee T, Delic D, Agyenim T, Calles C, Wagner R. Differential stringent control of Escherichia coli rRNA promoters: effects of ppGpp, DksA and the initiating nucleotides. Microbiology (Reading). 2011; 157: 2871–2879. doi:10.1099/mic.0.052357-0 PMID:21798983

10. Dalebroux ZD, Svensson SL, Gaynor EC, Swanson MS. ppGpp conjures bacterial virulence. Microbiol Mol Biol Rev. 2010;74: 171–199. doi:10.1128/MMBR.00046-09 PMID:20508246

11. Parshin A, Shiver AL, Lee J, Ozerova M, Schneidman-Duhovny D, Gross CA, et al. DksA regulates RNA polymerase in Escherichia coli through a network of interactions in the secondary channel that includes Sequence Insertion 1. Proc Natl Acad Sci U S A. 2015;112: E6862–6871. doi:10.1073/pnas.1521365112 PMID:26604313

12. Perron K, Comte R, van Delden C. DksA represses ribosomal gene transcription in Pseudomonas aeruginosa by interacting with RNA polymerase on ribosomal promoters. Mol Microbiol. 2005;56: 1087–1102. doi:10.1111/j.1365-2958.2005.04597.x PMID:15853892

13. Min KB, Yoon SS. Transcriptome analysis reveals that the RNA polymerase-binding protein DksA1 has pleiotropic functions in Pseudomonas aeruginosa. J Biol Chem. 2020;295: 3851–3864. doi:10.1074/jbc.RA119.011692 PMID:32047111

14. Gray MJ. Inorganic Polyphosphate Accumulation in Escherichia coli Is Regulated by DksA but Not by (p)ppGpp. J Bacteriol. 2019;201. doi:10.1128/JB.00664-18 PMID:30745375

15. Magnusson LU, Gummesson B, Joksimovic P, Farewell A, Nystrom T. Identical, independent, and opposing roles of ppGpp and DksA in Escherichia coli. J Bacteriol. 2007;189: 5193–5202. doi:10.1128/JB.00330-07 PMID:17496080

16. Pal RR, Bag S, Dasgupta S, Das B, Bhadra RK. Functional characterization of the stringent response regulatory gene dksA of Vibrio cholerae and its role in modulation of virulence phenotypes. J Bacteriol. 2012; 194: 5638–5648. doi:10.1128/JB.00518-12 PMID:22904284

17. Azriel S, Goren A, Rahav G, Gal-Mor O. The Stringent Response Regulator DksA Is Required for Salmonella enterica Serovar Typhimurium Growth in Minimal Medium, Motility, Biofilm Formation, and Intestinal Colonization. Infect Immun. 2016;84: 375–384. doi:10.1128/IAI.01135-15 PMID:26553464

18. Henard CA, Bourret TJ, Song M, Vazquez-Torres A. Control of redox balance by the stringent response regulatory protein promotes antioxidant defenses of Salmonella. J Biol Chem. 2010;285: 36785–36793. doi:10.1074/jbc.M110.160960 PMID:20851888

19. Blaby-Haas CE, Furman R, Rodionov DA, Artsimovitch I, de Crecy-Lagard V. Role of a Zn-independent DksA in Zn homeostasis and stringent response. Mol Microbiol. 2011;79: 700–715. doi:10.1111/j.1365-2958.2010.07475.x PMID:21255113

20. Stockert JC, Horobin RW, Colombo LL, Blazquez-Castro A. Tetrazolium salts and formazan products in Cell Biology: Viability assessment, fluorescence imaging, and labeling perspectives. Acta Histochem. 2018;120: 159–167. doi:10.1016/j.acthis.2018.02.005 PMID:29496266

21. Jiang Q, Chen J, Yang C, Yin Y Yao K. Quorum Sensing: A Prospective Therapeutic Target for Bacterial Diseases. Biomed Res Int. 2019;2019: 2015978. doi:10.1155/2019/2015978 PMID:31080810

22. Lee J, Zhang L. The hierarchy quorum sensing network in Pseudomonas aeruginosa. Protein Cell. 2015;6: 26–41. doi:10.1007/s13238-014-0100-x PMID:25249263

23. Lee KM, Yoon MY, Park Y, Lee JH, Yoon SS. Anaerobiosis-induced loss of cytotoxicity is due to inactivation of quorum sensing in Pseudomonas aeruginosa. Infect Immun. 2011;79: 2792–2800. doi:10.1128/IAI.01361-10 PMID:21555402

24. Nathwani D, Raman G, Sulham K, Gavaghan M, Menon V. Clinical and economic consequences of hospital-acquired resistant and multidrug-resistant Pseudomonas aeruginosa infections: a systematic review and meta-analysis. Antimicrob Resist Infect Control. 2014;3: 32. doi:10.1186/2047-2994-3-32 PMID:25371812

25. Yoon SS, Hennigan RF, Hilliard GM, Ochsner UA, Parvatiyar K, Kamani MC, et al. Pseudomonas aeruginosa anaerobic respiration in biofilms: relationships to cystic fibrosis pathogenesis. Dev Cell. 2002;3: 593–603. doi:10.1016/s1534-5807(02)00295-2 PMID:12408810

26. Arai H. Regulation and Function of Versatile Aerobic and Anaerobic Respiratory Metabolism in Pseudomonas aeruginosa. Front Microbiol. 2011;2: 103. doi:10.3389/fmicb.2011.00103 PMID:21833336

27. Hirai T, Osamura T, Ishii M, Arai H. Expression of multiple cbb3 cytochrome c oxidase isoforms by combinations of multiple isosubunits in Pseudomonas aeruginosa. Proc Natl Acad Sci USA. 2016;113: 12815–12819. doi:10.1073/pnas.1613308113 PMID:27791152

28. Ekici S, Yang H, Koch HG, Daldal F. Novel transporter required for biogenesis of cbb3-type cytochrome c oxidase in Rhodobacter capsulatus. mBio. 2012;3. doi:10.1128/mBio.00293-11 PMID:22294680

29. Choi KH, Schweizer HP. mini-Tn7 insertion in bacteria with single attTn7 sites: example Pseudomonas aeruginosa. Nat Protoc. 2006;1: 153–161. doi:10.1038/nprot.2006.24 PMID:17406227

30. Kim JS, Liu L, Fitzsimmons LF, Wang Y, Crawford MA, Mastrogiovanni M, et al. DksA-DnaJ redox interactions provide a signal for the activation of bacterial RNA polymerase. Proc Natl Acad Sci U S A. 2018;115: E11780–E11789. doi:10.1073/pnas.1813572115 PMID:30429329

31. Berger R, Resnati G, Metrangolo P, Weber E, Hulliger J. Organic fluorine compounds: a great opportunity for enhanced materials properties. Chem Soc Rev. 2011;40: 3496–3508. doi:10.1039/c0cs00221f PMID:21448484

32. Ikeda H, Yukawa M, Niiya T. Ab initio molecular orbital study of the reactivity of active alkyl groups. VII. Solvent effects on the formation of enolate isomers from 2-butanone with methoxide anion in methanol. Chem Pharm Bull (Tokyo). 2006;54: 731–734. doi:10.1248/cpb.54.731 PMID:16651780

33. Cloutier JF, Drouin R, Weinfeld M, O’Connor TR, Castonguay A. Characterization and mapping of DNA damage induced by reactive metabolites of 4-(methylnitrosamino)-1-(3-pyridyl)-1-butanone (NNK) at nucleotide resolution in human genomic DNA. J Mol Biol. 2001;313: 539–557. doi:10.1006/jmbi.2001.4997 PMID:11676538

34. Kim HY, Go J, Lee KM, Oh YT, Yoon SS. Guanosine tetra- and pentaphosphate increase antibiotic tolerance by reducing reactive oxygen species production in Vibrio cholerae. J Biol Chem. 2018;293: 5679–5694. doi:10.1074/jbc.RA117.000383 PMID:29475943

35. Klockgether J, Tummler B. Recent advances in understanding Pseudomonas aeruginosa as a pathogen. F1000Res. 2017;6: 1261. doi:10.12688/f1000research.10506.1 PMID:28794863

36. Paczkowski JE, Mukherjee S, McCready AR, Cong JP, Aquino CJ, Kim H, et al. Flavonoids Suppress Pseudomonas aeruginosa Virulence through Allosteric Inhibition of Quorum-sensing Receptors. J Biol Chem. 2017;292: 4064–4076. doi:10.1074/jbc.M116.770552 PMID:28119451

37. Yang YX, Xu ZH, Zhang YQ, Tian J, Weng LX, Wang LH. A new quorum-sensing inhibitor attenuates virulence and decreases antibiotic resistance in Pseudomonas aeruginosa. J Microbiol. 2012;50: 987–993. doi:10.1007/s12275-012-2149-7 PMID:23274985

38. Fan X, Liang M, Wang L, Chen R, Li H, Liu X. Aii810, a Novel Cold-Adapted N-Acylhomoserine Lactonase Discovered in a Metagenome, Can Strongly Attenuate Pseudomonas aeruginosa Virulence Factors and Biofilm Formation. Front Microbiol. 2017;8: 1950. doi:10.3389/fmicb.2017.01950 PMID:29067011

39. Guendouze A, Plener L, Bzdrenga J, Jacquet P, Remy B, Elias M, et al. Effect of Quorum Quenching Lactonase in Clinical Isolates of Pseudomonas aeruginosa and Comparison with Quorum Sensing Inhibitors. Front Microbiol. 2017;8: 227. doi:10.3389/fmicb.2017.00227 PMID:28261183

40. O’Loughlin CT, Miller LC, Siryaporn A, Drescher K, Semmelhack MF, Bassler BL. A quorum-sensing inhibitor blocks Pseudomonas aeruginosa virulence and biofilm formation. Proc Natl Acad Sci U S A. 2013; 110: 17981–17986. doi:10.1073/pnas.1316981110 PMID:24143808

41. Geske GD, O’Neill JC, Miller DM, Mattmann ME, Blackwell HE. Modulation of bacterial quorum sensing with synthetic ligands: systematic evaluation of N-acylated homoserine lactones in multiple species and new insights into their mechanisms of action. J Am Chem Soc. 2007;129: 13613–13625. doi:10.1021/ja074135h PMID:17927181

42. Pearson JP, Pesci EC, Iglewski BH. Roles of Pseudomonas aeruginosa las and rhl quorum-sensing systems in control of elastase and rhamnolipid biosynthesis genes. J Bacteriol. 1997; 179: 5756–5767. doi:10.1128/jb.179.18.5756-5767.1997 PMID:9294432

43. Pearson JP, Gray KM, Passador L, Tucker KD, Eberhard A, Iglewski BH, et al. Structure of the autoinducer required for expression of Pseudomonas aeruginosa virulence genes. Proc Natl Acad Sci U S A. 1994;91: 197–201. doi:10.1073/pnas.91.1.197 PMID:8278364

44. Steindler L, Venturi V. Detection of quorum-sensing N-acyl homoserine lactone signal molecules by bacterial biosensors. FEMS Microbiol Lett. 2007;266: 1–9. doi:10.1111/j.1574-6968.2006.00501.x PMID:17233715

45. Gentleman RC, Carey VJ, Bates DM, Bolstad B, Dettling M, Dudoit S, et al. Bioconductor: open software development for computational biology and bioinformatics. Genome Biol. 2004;5: R80. doi:10.1186/gb-2004-5-10-r80 PMID:15461798

46. Lee K, Lee KM, Go J, Ryu JC, Ryu JH, Yoon SS. The ferrichrome receptor A as a new target for Pseudomonas aeruginosa virulence attenuation. FEMS Microbiol Lett. 2016;363: fnw104. doi:10.1093/femsle/fnw104 PMID:27190289

47. Filloux A, Ramos JL. Preface. Pseudomonas methods and protocols. Methods Mol Biol. 2014;1149: v. doi:10.1007/978-1-4939-0473-0 PMID:24936603

