## Supplementary figures and tables for "Chemical Inhibitors of DksA1, a Conserved Bacterial Transcriptional Regulator, Suppressed Quorum Sensing-Mediated Virulence in *Pseudomonas aeruginosa*"

### Slide 1
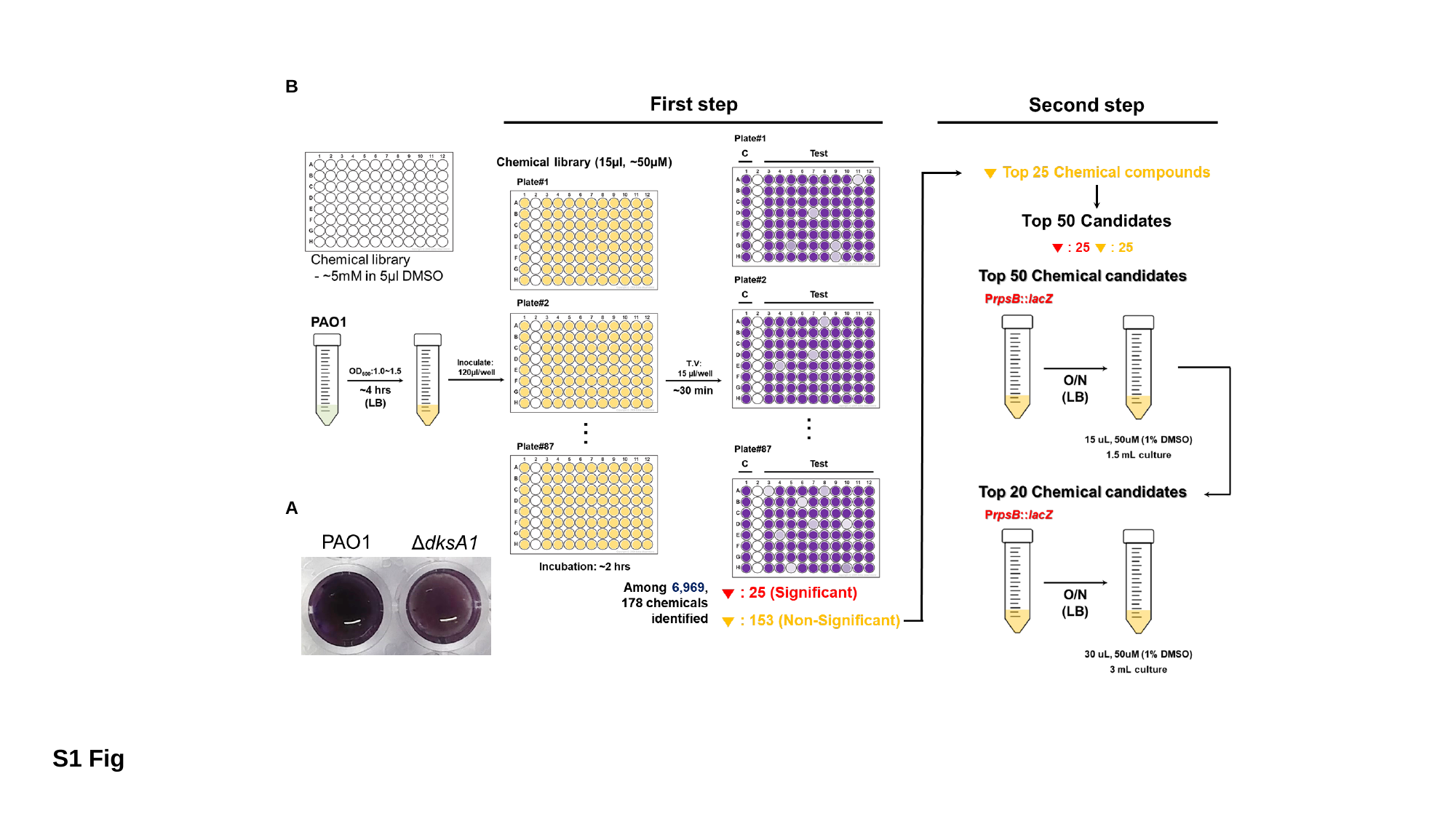

B
A
S1 Fig

### Slide 2
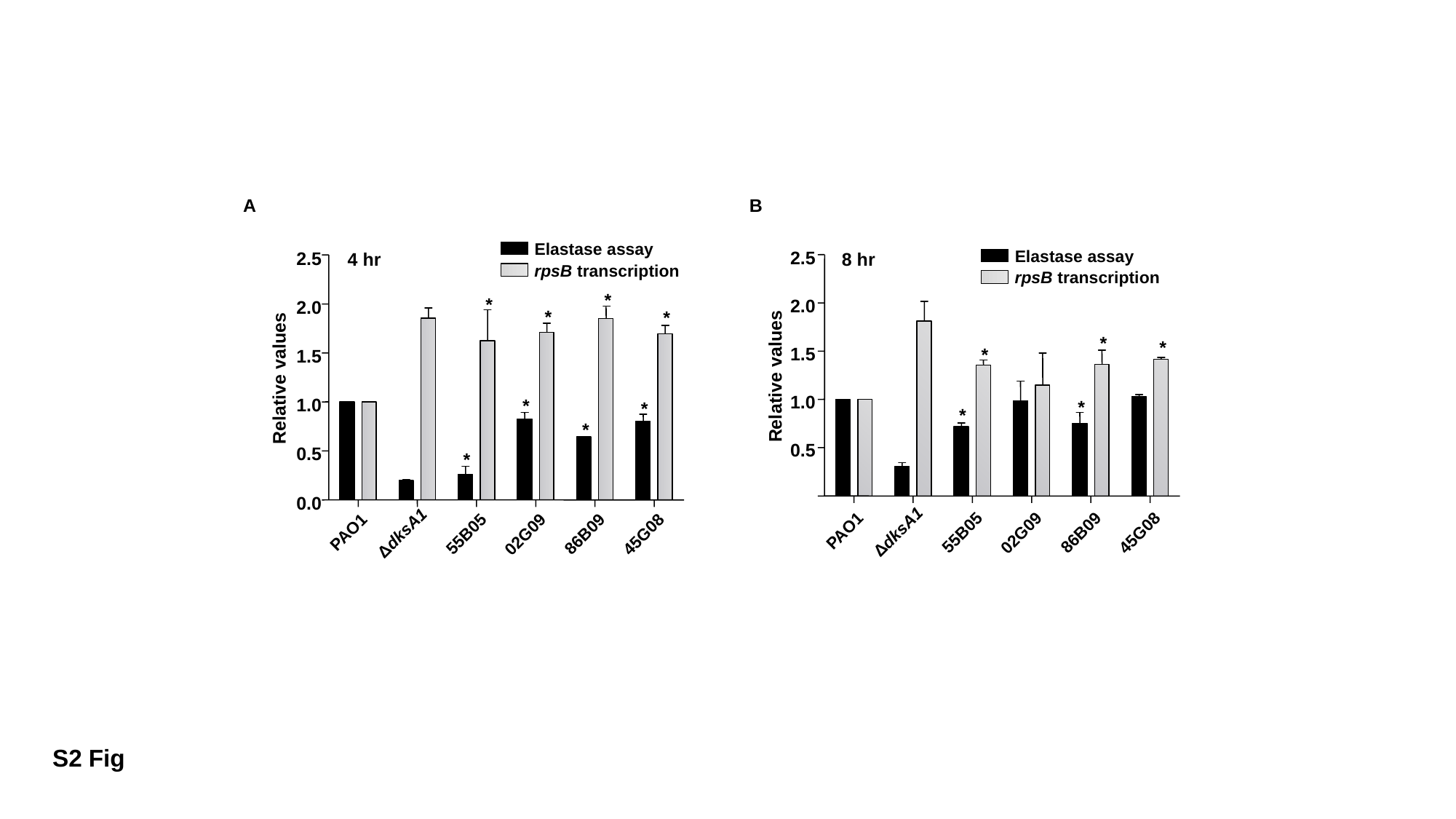

A
B
Elastase assay
4 hr
2.5
rpsB transcription
*
*
2.0
*
*
1.5
Relative values
*
*
1.0
*
0.5
*
0.0
PAO1
ΔdksA1
55B05
86B09
02G09
45G08
8 hr
Elastase assay
2.5
rpsB transcription
2.0
*
*
*
1.5
Relative values
*
1.0
*
0.5
PAO1
ΔdksA1
55B05
86B09
02G09
45G08
S2 Fig

### Slide 3
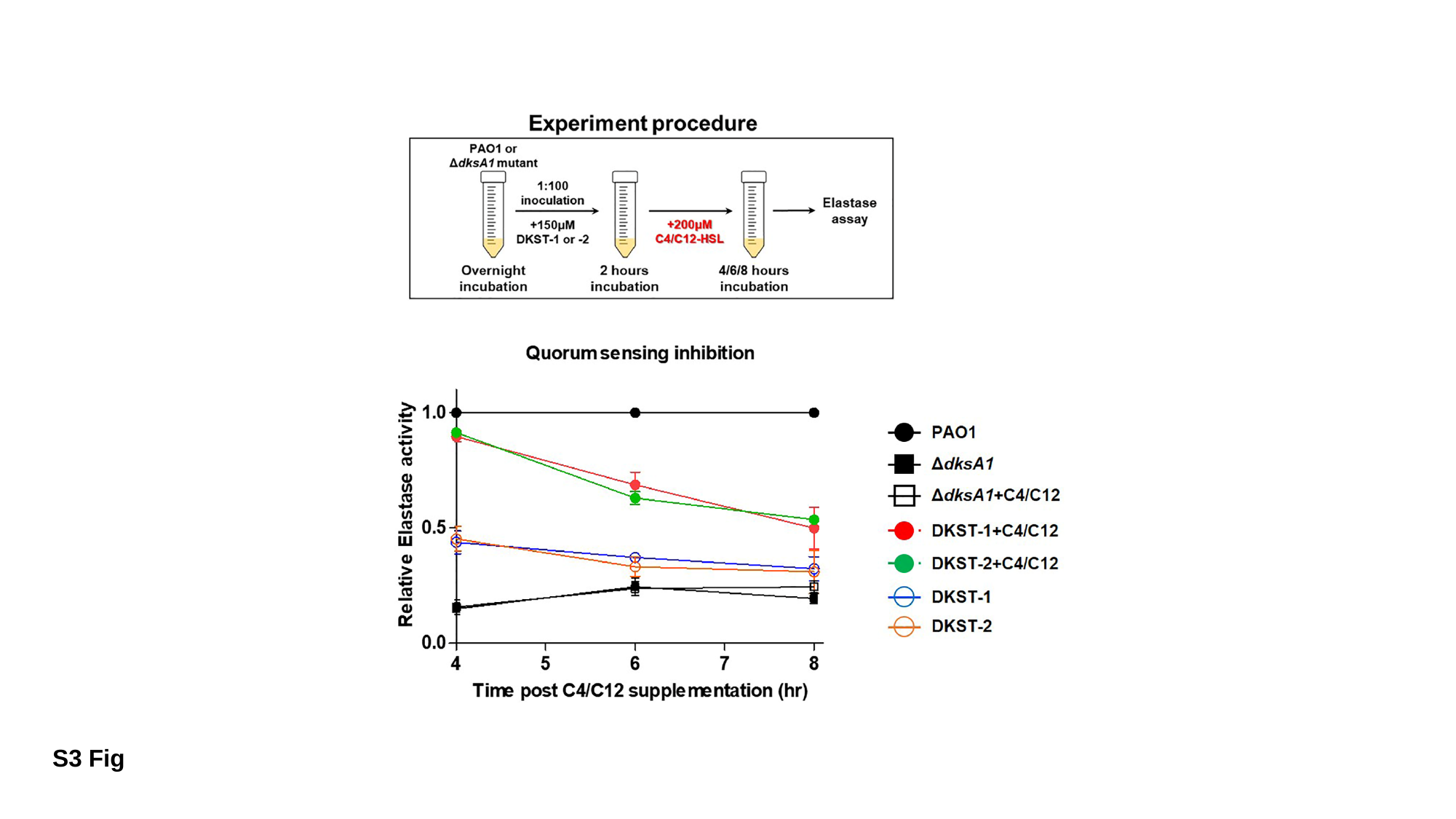

S3 Fig

### Slide 4
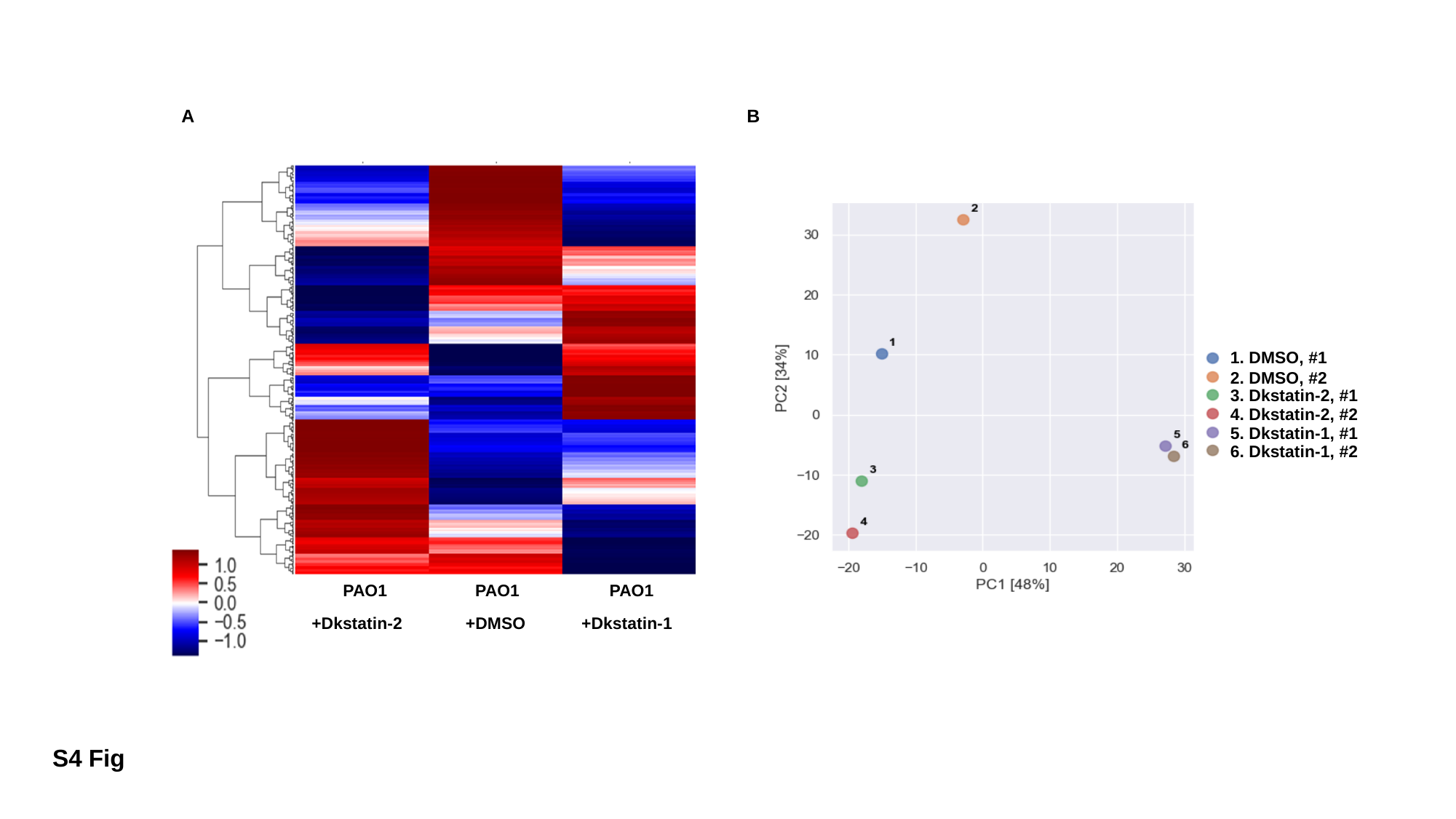

A
B
PAO1
PAO1
PAO1
+Dkstatin-2
+DMSO
+Dkstatin-1
1. DMSO, #1
2. DMSO, #2
3. Dkstatin-2, #1
4. Dkstatin-2, #2
5. Dkstatin-1, #1
6. Dkstatin-1, #2
S4 Fig

### Slide 5
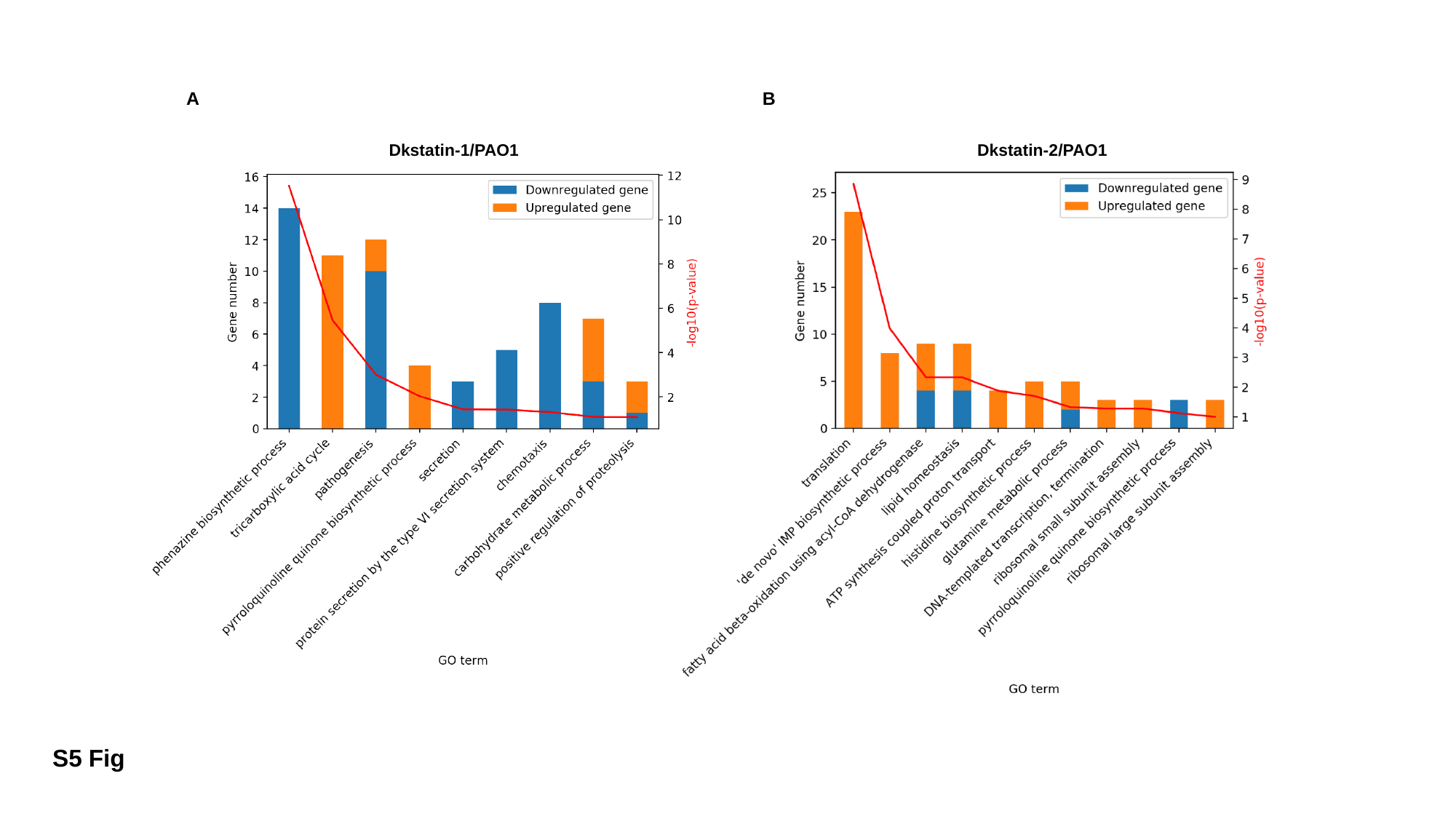

A
B
Dkstatin-1/PAO1
Dkstatin-2/PAO1
S5 Fig

### Slide 6
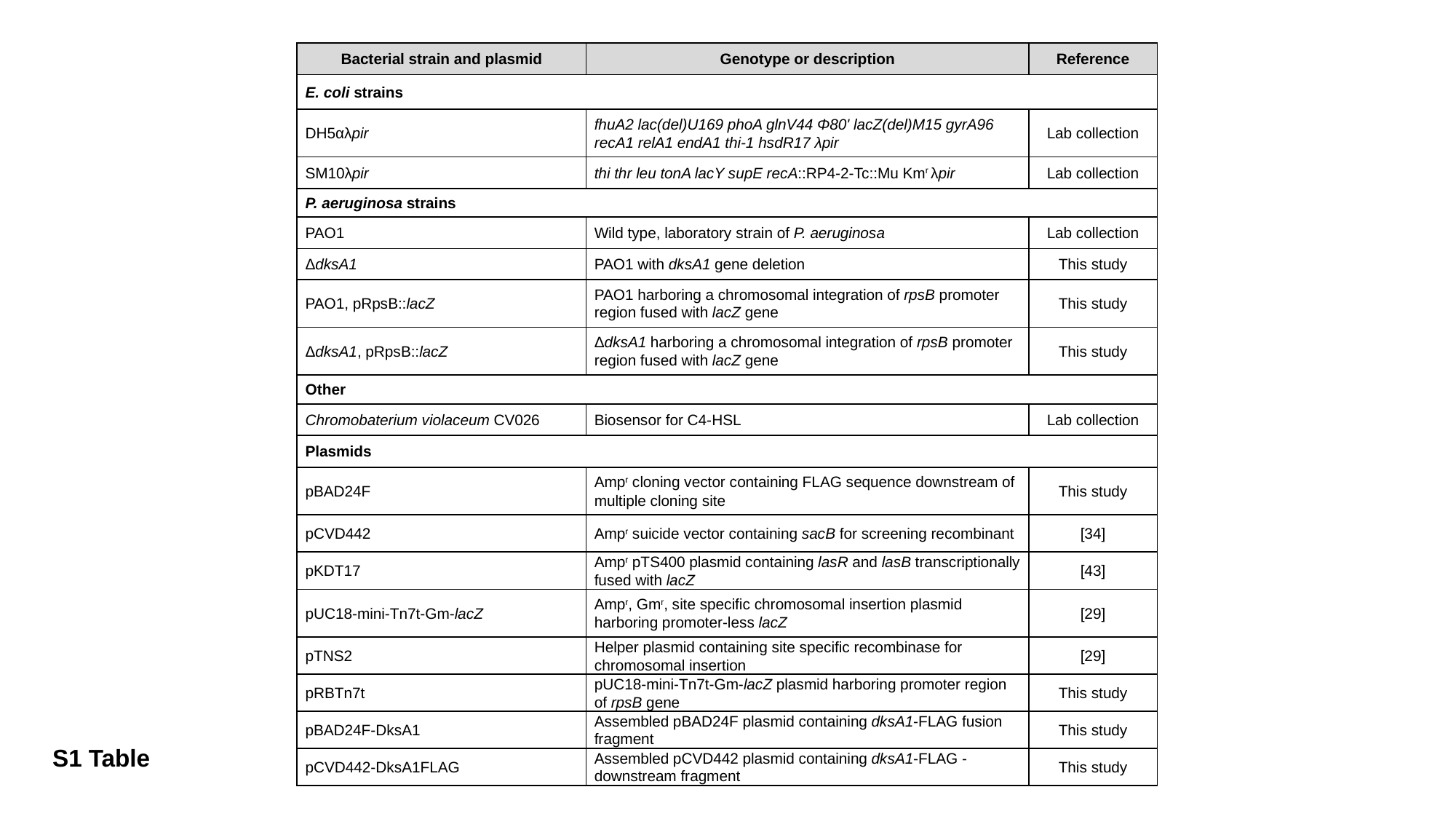

| Bacterial strain and plasmid | Genotype or description | Reference |
| --- | --- | --- |
| E. coli strains | | |
| DH5αλpir | fhuA2 lac(del)U169 phoA glnV44 Φ80' lacZ(del)M15 gyrA96 recA1 relA1 endA1 thi-1 hsdR17 λpir | Lab collection |
| SM10λpir | thi thr leu tonA lacY supE recA::RP4-2-Tc::Mu Km^r^ λpir | Lab collection |
| P. aeruginosa strains | | |
| PAO1 | Wild type, laboratory strain of P. aeruginosa | Lab collection |
| ΔdksA1 | PAO1 with dksA1 gene deletion | This study |
| PAO1, pRpsB::lacZ | PAO1 harboring a chromosomal integration of rpsB promoter region fused with lacZ gene | This study |
| ΔdksA1, pRpsB::lacZ | ΔdksA1 harboring a chromosomal integration of rpsB promoter region fused with lacZ gene | This study |
| Other | | |
| Chromobaterium violaceum CV026 | Biosensor for C4-HSL | Lab collection |
| Plasmids | | |
| pBAD24F | Amp^r^ cloning vector containing FLAG sequence downstream of multiple cloning site | This study |
| pCVD442 | Amp^r^ suicide vector containing sacB for screening recombinant | [34] |
| pKDT17 | Amp^r^ pTS400 plasmid containing lasR and lasB transcriptionally fused with lacZ | [43] |
| pUC18-mini-Tn7t-Gm-lacZ | Amp^r^, Gm^r^, site specific chromosomal insertion plasmid harboring promoter-less lacZ | [29] |
| pTNS2 | Helper plasmid containing site specific recombinase for chromosomal insertion | [29] |
| pRBTn7t | pUC18-mini-Tn7t-Gm-lacZ plasmid harboring promoter region of rpsB gene | This study |
| pBAD24F-DksA1 | Assembled pBAD24F plasmid containing dksA1-FLAG fusion fragment | This study |
| pCVD442-DksA1FLAG | Assembled pCVD442 plasmid containing dksA1-FLAG -downstream fragment | This study |
S1 Table

### Slide 7
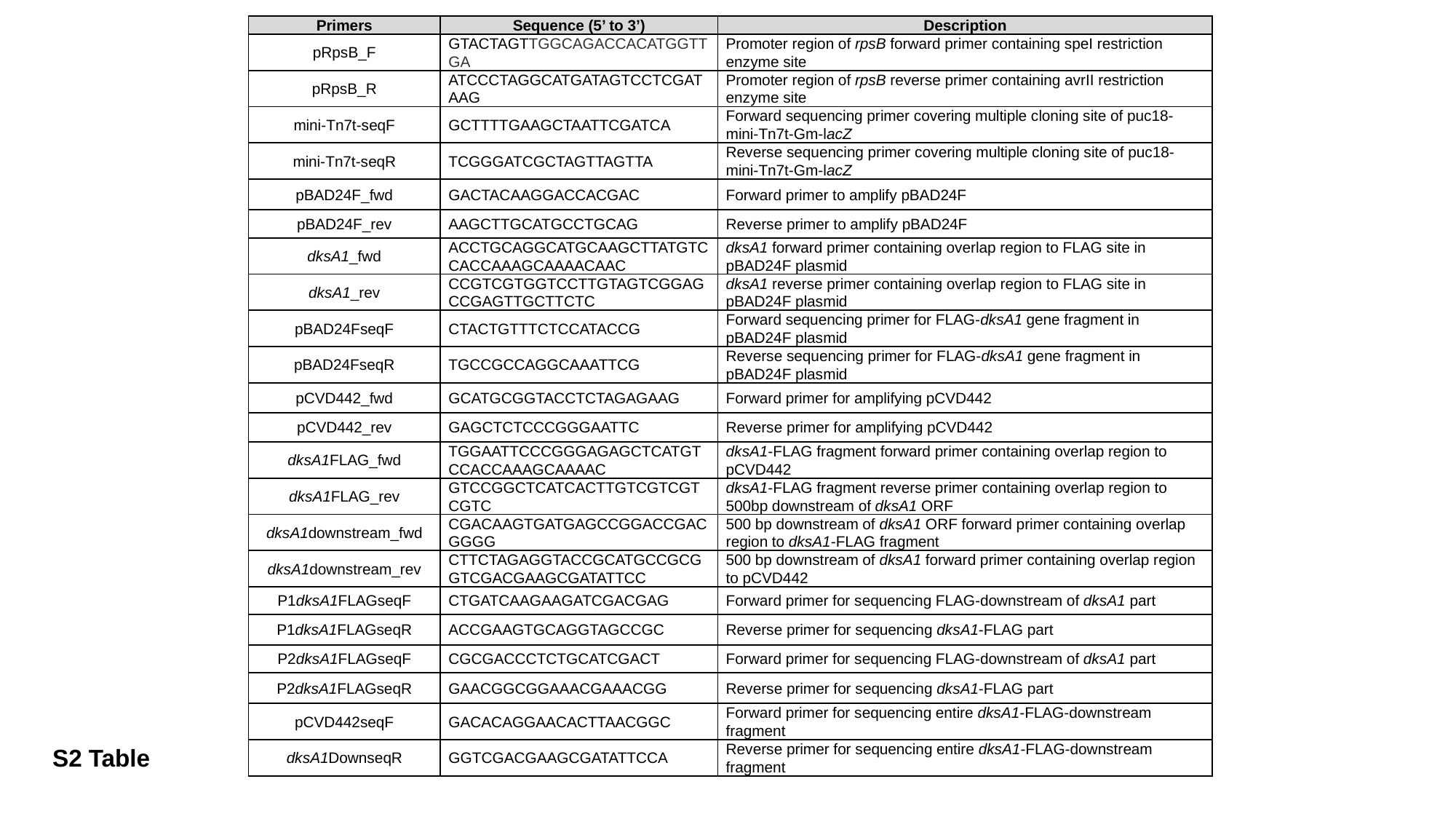

| Primers | Sequence (5’ to 3’) | Description |
| --- | --- | --- |
| pRpsB\_F | GTACTAGTTGGCAGACCACATGGTTGA | Promoter region of rpsB forward primer containing speI restriction enzyme site |
| pRpsB\_R | ATCCCTAGGCATGATAGTCCTCGATAAG | Promoter region of rpsB reverse primer containing avrII restriction enzyme site |
| mini-Tn7t-seqF | GCTTTTGAAGCTAATTCGATCA | Forward sequencing primer covering multiple cloning site of puc18-mini-Tn7t-Gm-lacZ |
| mini-Tn7t-seqR | TCGGGATCGCTAGTTAGTTA | Reverse sequencing primer covering multiple cloning site of puc18-mini-Tn7t-Gm-lacZ |
| pBAD24F\_fwd | GACTACAAGGACCACGAC | Forward primer to amplify pBAD24F |
| pBAD24F\_rev | AAGCTTGCATGCCTGCAG | Reverse primer to amplify pBAD24F |
| dksA1\_fwd | ACCTGCAGGCATGCAAGCTTATGTCCACCAAAGCAAAACAAC | dksA1 forward primer containing overlap region to FLAG site in pBAD24F plasmid |
| dksA1\_rev | CCGTCGTGGTCCTTGTAGTCGGAGCCGAGTTGCTTCTC | dksA1 reverse primer containing overlap region to FLAG site in pBAD24F plasmid |
| pBAD24FseqF | CTACTGTTTCTCCATACCG | Forward sequencing primer for FLAG-dksA1 gene fragment in pBAD24F plasmid |
| pBAD24FseqR | TGCCGCCAGGCAAATTCG | Reverse sequencing primer for FLAG-dksA1 gene fragment in pBAD24F plasmid |
| pCVD442\_fwd | GCATGCGGTACCTCTAGAGAAG | Forward primer for amplifying pCVD442 |
| pCVD442\_rev | GAGCTCTCCCGGGAATTC | Reverse primer for amplifying pCVD442 |
| dksA1FLAG\_fwd | TGGAATTCCCGGGAGAGCTCATGTCCACCAAAGCAAAAC | dksA1-FLAG fragment forward primer containing overlap region to pCVD442 |
| dksA1FLAG\_rev | GTCCGGCTCATCACTTGTCGTCGTCGTC | dksA1-FLAG fragment reverse primer containing overlap region to 500bp downstream of dksA1 ORF |
| dksA1downstream\_fwd | CGACAAGTGATGAGCCGGACCGACGGGG | 500 bp downstream of dksA1 ORF forward primer containing overlap region to dksA1-FLAG fragment |
| dksA1downstream\_rev | CTTCTAGAGGTACCGCATGCCGCGGTCGACGAAGCGATATTCC | 500 bp downstream of dksA1 forward primer containing overlap region to pCVD442 |
| P1dksA1FLAGseqF | CTGATCAAGAAGATCGACGAG | Forward primer for sequencing FLAG-downstream of dksA1 part |
| P1dksA1FLAGseqR | ACCGAAGTGCAGGTAGCCGC | Reverse primer for sequencing dksA1-FLAG part |
| P2dksA1FLAGseqF | CGCGACCCTCTGCATCGACT | Forward primer for sequencing FLAG-downstream of dksA1 part |
| P2dksA1FLAGseqR | GAACGGCGGAAACGAAACGG | Reverse primer for sequencing dksA1-FLAG part |
| pCVD442seqF | GACACAGGAACACTTAACGGC | Forward primer for sequencing entire dksA1-FLAG-downstream fragment |
| dksA1DownseqR | GGTCGACGAAGCGATATTCCA | Reverse primer for sequencing entire dksA1-FLAG-downstream fragment |
S2 Table
